# Mapping a *Toxoplasma gondii* interactome by crosslinking mass spectrometry and machine learning

**DOI:** 10.1101/2025.04.28.651050

**Authors:** Tadakimi Tomita, Elizabeth Weyer, Rebekah Guevara, Simone Sidoli, Jennifer T Aguilan, Louis M. Weiss

## Abstract

*Toxoplasma gondii,* a widespread human parasite, persists in hosts through complex molecular interactions. Protein-protein interactions (PPIs) underpin essential biological processes, including parasite-host interactions and cellular invasion. Herein, we utilized advanced crosslinking mass spectrometry (XL-MS) techniques to map a *T. gondii* tachyzoite cytosolic extract interactome. By integrating MS-cleavable and non-cleavable analysis, we identified a total of 405 unique PPIs at medium confidence (FDR < 5%) and 226 at high confidence (FDR < 1%), revealing both known and novel interactions within critical cellular complexes such as the ribosome, proteasome, and dense granule proteins. Structural validation confirmed spatial proximity of crosslinked residues, while comparative analyses against existing datasets (hyperLOPIT, ToxoNET, STRING) corroborated the biological relevance of identified interactions. Furthermore, we introduced a machine learning approach leveraging biological annotations and experimental data to significantly enhance the detection and validation of PPIs. Our findings not only provide a refined view of *T. gondii*’s molecular architecture, but also highlight the utility of XL-MS coupled with computational tools in dissecting complex parasite proteomes. The XL-MS interactome map provides a new valuable resource for understanding parasite biology and developing targeted therapeutic strategies.

## Introduction

*Toxoplasma gondii* is a prevalent human parasite that is estimated to chronically infect a quarter of the human population (1). HIV-related cerebral toxoplasmosis remains a significant problem in people with HIV infection with CD4+ counts under 200 cells/mm^3^ (2). While it is the most common cause of inflammatory mass lesions in the central nervous system in these patients, the availability of combination antiretroviral therapy (cART) has significantly reduced the incidence of cerebral toxoplasmosis in people living with HIV/AIDS (3). Despite the lower incidence rates, it remains an important problem and research is needed to develop new approaches to ameliorate the pathogenesis of this central nervous system illness.

Protein-protein interactions (PPIs) are vital for the understanding of the biologic process in a cell. These interactions allow proteins to bind together, creating structures and forming functional biochemical machines, which regulate protein function. Such processes are instrumental in orchestrating cellular activities, including replication, movement, and specifically in the case of *Toxoplasma gondii*, the infection of host cells. Therefore, cataloging proteome-wide PPIs extends beyond mere stamp collecting; it is a critical step in unraveling the intricate complexity of biological life.

PPIs are traditionally studied using methods such as affinity purification, co-immunoprecipitation, BioID-based proximity labeling, and yeast-two-hybrid (Y2H) methods. Affinity purification-mass spectrometry (AP-MS) isolates specific proteins from complex mixtures via tags or antibodies, followed by mass spectrometric identification. While precise, AP-MS struggles to capture weak or transient PPIs and requires genetic modifications for affinity tags and specific antibodies, making proteome-level studies challenging. Biotin proximity labeling (BioID) employs a biotin ligase fused to a target protein to covalently attach biotin to nearby proteins that may interact with the target. These biotinylated proteins are subsequently isolated via streptavidin affinity purification. Although this method is effective in detecting weak and transient interactions, it does not confirm direct physical contact and still relies on genetic tagging, which may restrict its broader applicability. Emerging techniques like photoactivable unnatural amino acid crosslinking more directly explore interactions, but also demand genetic manipulation of each protein (4). Conversely, approaches such as hyperLOPIT (5) and co-elution (6) are used for large-scale mapping of subcellular locations and protein complexes in parasites, achieving proteome-wide insights, but with lower resolution concerning specific organelles and complexes. Crosslinking mass spectrometry (XL-MS) has recently evolved into a powerful tool for studying protein–protein interactions, due to advancements in peptide mass spectrometry and computational methods. The technique works by forming covalent bonds that effectively “freeze” the proteins in their native state. A chemical crosslinker is used to impose spatial constraints that mirror the in-solution conformations of intact proteins.

A new generation of enrichable MS-cleavable crosslinkers such as DSBSO (7) has greatly improved XL-MS. Click reactive groups have been incorporated to enrich crosslinked peptides from unmodified peptides successfully used for proteome-level studies. These include MS non-cleavable but more hydrolysis-resistant *N*-succinimidyl carbamate crosslinker NNP9 (8) and the MS-cleavable carboxyl-selective crosslinker BAP (9). Click enrichment of digested peptides rather than proteins increases the recovery of the labeled proteins (10). Bypassing biotin-streptavidin and Cu(I)-catalyzed azide-alkyne cycloaddition (CuAAC), through the simple use of strain-promoted azide-alkyne cycloaddition (SPAAC) by Dibenzocyclooctyne (DBCO) coupled beads, increases the enrichment efficiency (11). In addition, enrichment of larger peptides by size exclusion columns increases recovered crosslinks and reduces monolinks (12, 13). Use of high-pH reverse-phase fractionation further reduces the complexity of samples and expands coverage (13). On the instrument-side, the use of high-field asymmetric waveform ion mobility spectrometry (FAIMS) improves depth and sensitivity by enriching highly charged ions (12, 14). Moreover, to further optimize peptide identification, stepped higher-energy collisional dissociation (HCD) fragmentation can be implemented (15).

Herein we demonstrate that the XL-MS protocol we developed generated crosslinking data that is consistent with known protein-protein interactions and experimentally derived structures. Additionally, we developed and applied a machine learning approach to increase the number of PPI identified in our crosslinking data set. Finally, we demonstrate the utility of XL-MS data in predicting protein structure, both of individual proteins and of protein complexes.

## Methods

### Chemicals

Chemicals used in these experiments were purchased from commercial suppliers: e.g. Azide-A-DSBSO (Millipore Sigma: 909629), DBCO agarose beads (Vector laboratories), Trypsin Gold 100 ug (Promega), Lys-C (MedChemExpress), Protease inhibitor cocktail cOmplete EDTA free (Roche), and high pH reversed-phase peptide fractionation kit (Pierce). We observed significant variations in the quality of DSBSO across different lots from Millipore Sigma. To ensure efficient crosslinking and maintain consistency in experimental outcomes, we recommend conducting a preliminary crosslinking assay with each new lot prior to its use in large-scale experiments and/or validation of lots of DSBSO by NMR.

### Cell culture

An ME49 strain of *T. gondii* with deletions of the KU80 gene (TGME49_312510) and hypoxanthine-xanthine-guanine phosphoribosyl transferase (TGME49_200320), herein referred to as ME49Q (16) was cultured in human foreskin fibroblasts (HFF, ATCC: CRL-1634; Hs27) in Dulbecco’s modified Eagle medium supplemented with penicillin-streptomycin at 37°C 5% CO_2_. The experiment was performed as duplicates. Eight 150 mm plates of HFF per sample were infected with this ME49Q strain and cultured for three days. Parasites were lysed out from host cells by passing through a 27G needle three times and a 5 µm membrane filter once. Parasites were then washed three times with PBS followed by hypotonic lysis that was performed as described previously (17). Briefly, the cell pellet was resuspended with 100 µl of 10 mM HEPES pH 7.5 with cOmplete protease inhibitor cocktail EDTA free and incubated on ice for 10 minutes. The lysate was spun at 16,000 x g and the supernatant was harvested. Hypotonic extraction was repeated a total of five times. The soluble lysates were combined and then 10x PBS was added to the lysate at 1x final concentration.

### Crosslinking and Digestion

The crosslinking protocol was performed following a previously published method (17) with the following modifications. Lysates were adjusted to 1 µg/µL with 1x PBS. The crosslinker Azide-A-DSBSO was added to the lysate at a concentration of 5 mM This was followed by incubation for 2 hours at room temperature while the tube rotated at 10 rpm. Crosslinking was terminated by adding Tris pH 8 at 50 mM and incubation for 5 minutes at room temperature. Digestion was performed using the filter-aided sample preparation method (FASP) (18, 19) as follows: the samples were washed with 8 M urea in 25 mM ammonium bicarbonate three times using a 3kDa nominal cutoff Amicon Ultra filter device (Millipore Sigma). The proteins were reduced with Tris (2-carboyethyl) phosphine at 2 mM for 30 minutes and alkylated with iodoacetamide at 20 mM in the dark for 30 minutes. The samples were washed once with 8 M urea in 25 mM ammonium bicarbonate and digested in 6.4 M urea in 25 mM ammonium bicarbonate with 1:100 (w/w, protease:protein) LysC for 4 hours at 37°C. Subsequently, the sample was diluted with 25 mM ammonium bicarbonate to 1 M urea and incubated overnight at 37°C with Trypsin Gold at 1:50 w/w. After the digestion, acetonitrile was added to the final concentration of 10% and incubated with 40 µl of DBCO agarose beads at 4 °C overnight. The DBCO beads were washed three times with 1 M NaCl and 10% acetonitrile in 25 mM ammonium bicarbonate sequentially. The peptides were eluted with 10% trifluoroacetic acid (TFA) by rotating the tube at room temperature for 30 minutes at 5 rpm. The beads were eluted twice, and the eluates were neutralized with ammonium bicarbonate to yield a final concentration of 100 mM ammonium bicarbonate in the tube. The beads were then eluted with 50% acetonitrile in 25 mM ammonium bicarbonate twice. All eluates were then combined and vacuum dried. The peptides were fractionated with high pH reversed-phase peptide fractionation kit following the manufacturer’s protocol (Pierce). The elution was performed with 5, 10, 15, 20, 25, 30, 35, and 50% acetonitrile sequentially and two fractions were combined (5+25, 10+30, 15+35, and 20+50%) to make four fractions per sample.

### Crosslinked peptide enrichment by SEC

An additional sample was enriched using size exclusion chromatography (SEC) rather than high-pH fractionation. Following click chemistry-based purification, the dried peptides were dissolved in 30% acetonitrile (ACN) with 0.1% TFA to achieve a concentration of 20 µg/µl. Two hundred µg of peptides were then injected into a Superdex 30 Increase 3.2/300 column (Cytiva) and eluted isocratically with the same buffer. Fifty-µl fractions were collected, dried, and subsequently analyzed by mass spectrometry. The result of this fractionation is provided in Figure S1. In summary, the fraction from 1.18 mL to 1.23 mL exhibited the highest ratio of crosslinked to non-crosslinked species, while the fraction from 1.23 mL to 1.28 mL contained the greatest number of crosslinks.

### Mass spectrometric analysis

Samples were resuspended in 0.1% TFA and loaded onto a Dionex RSLC Ultimate 3000 (Thermo Scientific), coupled online with an Orbitrap Exploris 480 (Thermo Scientific) with FAIMS Pro attached. Chromatographic separation was performed with a two-column system, consisting of a C-18 trap cartridge (300 µm ID, 5 mm length) and a picofrit analytical column (75 µm ID, 25 cm length) packed in-house with reversed-phase Repro-Sil Pur C18-AQ 3 µm resin. Peptides were separated using a 180 minutes gradient from 4-30% buffer B (buffer A: 0.1% formic acid, buffer B: 80% acetonitrile + 0.1% formic acid) at a flow rate of 300 nl/min. The mass spectrometer was set to acquire spectra in a data-dependent acquisition (DDA) mode. Briefly, the full MS scan was set to 375-1600 m/z with a resolution of 120,000 (at 200 m/z). MS/MS was performed at 60,000 resolution and AGC target of 1×10e4 and stepped HCD normalized collision energy of 19, 25, 32% % (11, 15, 20) with FAIMS compensation voltages of −50, −60, and −75 (14).

The data was analyzed with Proteome Discoverer 2.5 with IMP-MS2 spectrum Processor for de-isotoping and MS Amanda 2.0 with the following parameters: maximum missed cleavage 4, MS1 tolerance 10 ppm, MS2 tolerance 20 ppm, static modification: carbamidomethyl 57.021 Da (C), dynamic modifications: oxidation 15.995 Da (M). Spectrum searches were performed to ToxoDB (toxodb.org) release 67 (21). Data sets were analyzed with MS Annika 2.0 (22) as MS-cleavable crosslinkers with the following parameters: DSBSO 308.039 Da (K), alkene 54.0156 Da, and thiol 236.0177 Da, additional crosslink doublet 200.0177 on K, S, T, and Y residues. Additionally, data were analyzed with pLink 2 (23) as non-cleavable crosslinker with the following parameters: DSBSO 308.378 Da (linker mass), 326.394 Da (mono mass), Precursor14 276.358 Da (linker mass), 294.374 Da (mono mass).

### Data Processing and Machine learning

The data generated by Annika 2.0 and pLink 2 were processed using R (24) script with tidyverse (25) packages. The hyperLOPIT (5), CRISPR fitness score (26), and GO term data were downloaded from ToxoDB release 67 (toxodb.org). The STRING data was downloaded from STRING version 12.0 (string-db.org), Uniprot names were converted to ToxoDB ID using UniProt ID mapping tool. Machine learning was performed on Google Colab with LightGBM (27). Hyperparameter tuning was performed with Optuna (28). The Python and R scripts used in this study are in the supplementary file (Supplemental file 1).

### Data visualization

Interactomes were visualized with Cytoscape 3 (29). Interactomes with residue-to-residue interactions were visualized using xiView (30). Structural mapping to the known structures and measurement of crosslink distance were performed with ChimeraX (31) with XMAS plugin (32). Distance of random K-K residues were measured using PyMOL and python script.

### Data Deposition

Proteomic datasets were deposited to ProteomeXhange under the project accession number PXD062479 and to ToxoDB (EuPATHdB).

## Results and Discussions

### Determining the cytoplasmic crosslinking interactome

To establish a *Toxoplasma* interactome using XL-MS, we crosslinked tachyzoite soluble cytosolic lysate. Tachyzoites were selected as the starting material since they can be produced in large quantities and readily used for crosslinking. A schematic representation of the experiment is depicted in Figure 1A. Using the cleavable detection method, the XLMS analysis generated a total of 18692 crosslinking spectrum matches and 3551 unique residue-residue pairs, including 3045 intra-protein links and 506 inter-protein links, and 305 PPI at medium confidence (FDR < 5%) from the biological triplicates (Table 1). To get the overall composition of proteins in the sample, gene ontology analysis was performed on all 305 proteins that were crosslinked. Highly significant terms include ribosome, proteasome complex, cytoplasm, and translation initiation complex representing a wide variety of intracellular soluble tachyzoite lysates (Table 2). To increase the number of crosslinked peptides, the same data were also searched for non-MS-cleavable crosslinks, despite using DSBSO, which is designed for the MS-cleavable protocol. This approach is based on the rationale that some crosslinked peptides may not dissociate at the crosslinker during mass spectrometry analysis, but these peptides can still fragment allowing for their identification. Although the non-cleavable search yielded slightly fewer results compared to the cleavable search, a substantial number of additional spectra were identified at false discovery rates (FDR) of <1% and <5% (Table 1). This led to a 54–92% increase in residue-residue interactions (RRIs), demonstrating the benefits of this method. Additionally, only one-third of protein-protein interactions (PPIs) were shared between the cleavable and non-cleavable searches, underscoring the advantage of employing both search strategies to achieve broader coverage. Overall, the identification of PPIs improved substantially, when we combined the detection methods, with a total of 391 PPIs identified at an FDR <5% and 239 PPIs at an FDR <1%.

**FIGURE 1:**
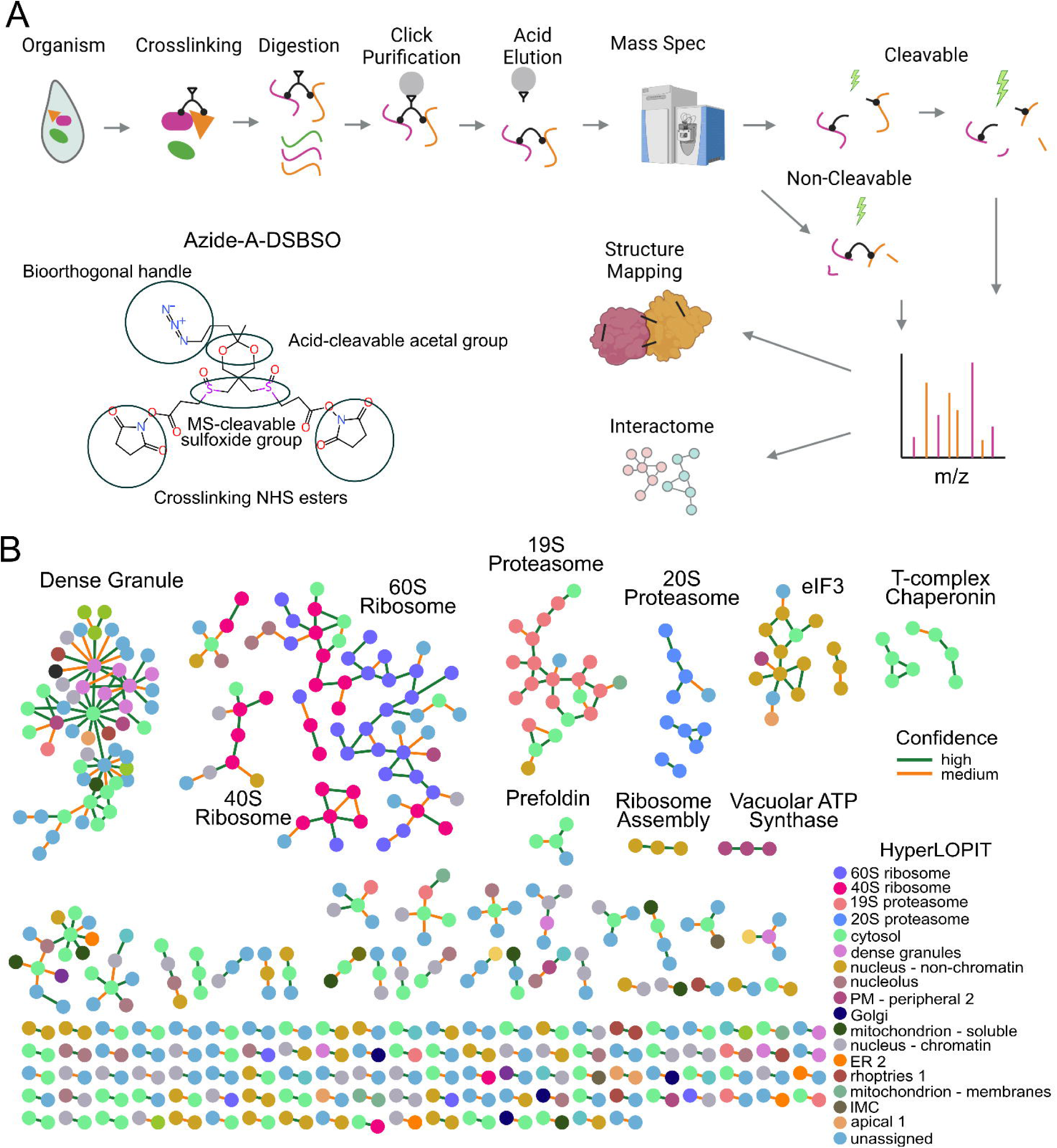
Cross Linking Mass Spectrometry. **(A)** The schematic illustrates the overall process of crosslinking mass spectrometry used in this study. The chemical structure shown is that of azide-a-DSBSO, the crosslinker, which contains various functional groups highlighted by circles. **(B)** Interactome networks derived from crosslinking mass spectrometry analyses of T. gondii cytosolic proteins. Each node represents a protein and is color-coded according to its subcellular localization, as determined by hyperLOPIT assignments. The edges are color-coded based on the confidence of identification (high: FDR < 1%; medium: 1% < FDR < 5%).

### Evaluating protein-protein interactions

Our interactome analysis generated networks with 405 nodes and 527 edges at FDR of < 5%, and 226 nodes and 304 edges at FDR of < 1%, respectively. Figure 1B displays the networks with unique protein-protein interactions and genes are colored based on hyperLOPIT localization assignments. There are several distinct networks of the interactome. As expected for cytosolic proteome, the most prominent complexes are the abundant 60S ribosome and 40S ribosome complexes comprising more than 50 proteins. The 19S and 20S proteasome complexes form two separate networks, reflecting how the proteasomes make this structure. The eukaryotic initiation factor 3 (eIF3) complex is also detected. Strikingly, seven dense granular proteins form a densely connected subnetwork that has connection to SRS protein network. This indicates that the crosslinking MS can be used for proteins enclosed in membranous secretory vesicles inside the cytosol due to the membrane permeability of the crosslinker.

To validate the feasibility of the interactomes that were generated by the crosslinking data, we compared our findings with other independently assessed large-scale proteomics data, namely subcellular localization by hyperLOPIT (5) and protein-complexes by co-elution ToxoNET (33). Of all pairs found in independent dataset, excluding missing values, 72.0% of protein-protein interactions are in the same location in hyperLOPIT and 62.1% are in the same cluster in ToxoNET (Table 3). The high rate of agreement, relative to disagreement and in comparison to random pairs, indicates that these protein pairs are likely to represent genuine interactions. We further evaluated the concordance among additional biological features—such as Gene Ontology (GO) components, functions, processes, EC numbers, Pfam IDs, and STRING pairs— and observed a similar trend: high-confidence hits exhibited greater agreement, while lower-confidence hits showed reduced concordance.

### Improving crosslinking using machine learning

A false discovery rate (FDR) threshold of less than 1% is routinely applied to classify mass spectra as “high confidence,” and 5% is commonly used as “medium confidence” ensuring the reliability of spectral quality. However, an analysis of the histogram depicting spectral hits matched to both the target and decoy (reversed sequence) databases reveals that a substantial proportion of hits within higher FDR ranges are likely true positives. Specifically, 86% of medium-confidence hits (1% < FDR < 5%) and 64% of low-confidence hits (5% < FDR < 10%) are expected to be theoretically genuine (Figure 2A). Further investigation of these medium- and low-confidence PPI has enabled the identification of highly probable crosslinks, including the proteins that are known to be in the same complex, such as numerous ribosomal proteins, proteasomal proteins, elongation factor 1 complex (EF1), eukaryotic initiation factor 3 complex (eIF3), and prefoldin complex. A higher agreement percentage among the high-confidence hits (Table 3), compared to those of low or random pairs, suggests that we can leverage our prior biological knowledge to effectively distinguish potential true hits from false ones. Consequently, we attempted to salvage these middle to low-score hits by applying biological features through machine learning algorithms (Figure 2B). We selected readily available biological features from ToxoDB, which include tachyzoite transcriptomic data, CRISPR fitness scores, hyperLOPIT, Gene Ontology, PFam, and EC numbers. Additionally, we incorporated interactions from the STRING database and experimentally generated ToxoNET data.

**FIGURE 2:**
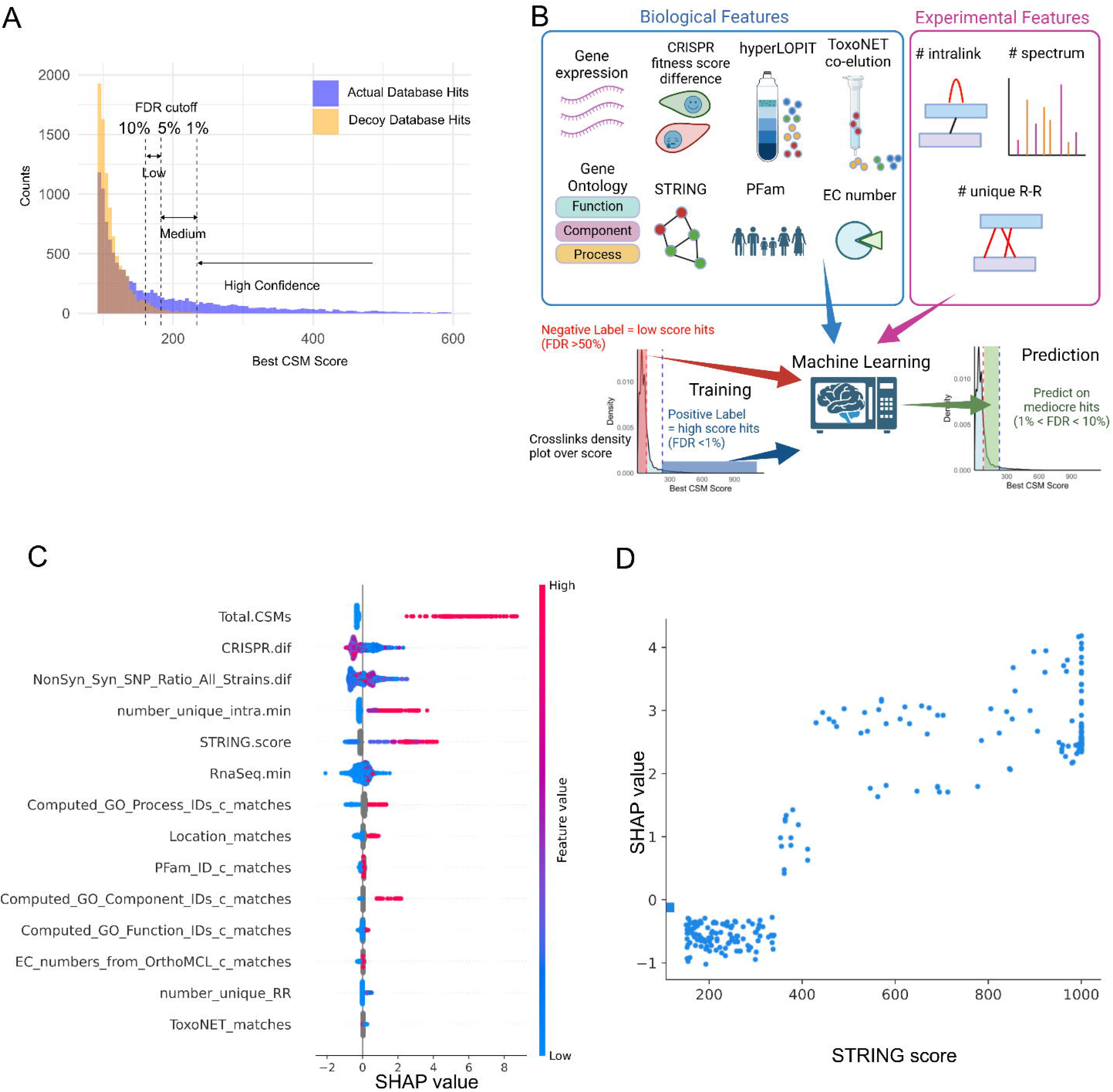
Machine Learning. (**A**) Histogram of identification score (best CSM score) in true hits (blue) and decoy hits (yellow). The histogram illustrates the theoretical true hits within the medium and low confidence regions, with the confidence cutoff thresholds indicated by dotted lines. The blue areas in the low and medium confidence regions, located above the overlap with decoy hits, represent the theoretical true hits that can be salvaged. (**B**) Schematic illustrating the recovery of medium- to low-confidence hits using machine learning based on biological and experimental features. (**C**) Analysis of feature importance and individual prediction contributions in the model. SHapley Additive exPlanations (SHAP) values are presented, with positive values indicating that the feature increases the prediction and negative values suggesting a decrease. Features are ranked from most to least influential. Feature values are color-scaled from high (red) to low (blue), with missing values shown in gray. (**D**) Relationship between STRING score and feature importance. Higher STRING scores contribute to positive predictions (as indicated by SHAP values), whereas lower scores contribute to negative predictions in the model construction.

In addition to the biological features, recent publications have identified ways to improve crosslink identification, taking into account crosslinks based on the presence of corresponding proteins detected in intralinks (intra-type dependent), and the presence of multiple unique residue-residue interlink pairs (inter-dependent) (34, 35). We therefore included similar features that can be easily implemented for machine learning. Those features are the number of unique residue-residue pairs in the PPI and the number of intralink spectra (Figure 2B).

We chose a gradient-boosting decision-tree algorithm LightGBM (27) because of its ability to use tabulated data containing both categorical and numerical features, as well as its native support for missing values in binary classification tasks. High-confidence pairs (FDR < 1%) were used as the positive training data, while the lowest-confidence pairs (FDR > 50%) served as negative training data. The resulting model was subsequently evaluated on an independent test set, achieving an accuracy of 0.987 and a Receiver Operating Characteristic Area Under the Curve (ROC AUC) of 0.959. One of the key advantages of a decision-tree-based algorithm is the interpretability of the features used in model construction. SHapley Additive exPlanation (SHAP) (36) were employed to quantify the contribution of each feature to the model’s predictions. Not surprisingly, the total number of CSM emerged as the most influential feature, with higher counts strongly predicting true hits (Figure 2C). Additionally, the number of unique intralinks (with a minimum of two) was found to be a strong predictor of interaction, which is consistent with recent publications (35). In contrast, the number of unique residue-to-residue interactions did not contribute significantly to the model – likely because most interactions were represented by a single hit, resulting in insufficient variability for effective prediction.

Biological features also contributed significantly to the construction of the predictive model. One of the most influential biological features was the difference in the CRISPR fitness scores between protein pairs; proteins with similar fitness scores were more likely to form true crosslinks. This observation suggests that, on a large proteomic scale, essential genes tend to form complexes that enhance cellular fitness, whereas dispensable genes tend to form complexes associated with non-essential functions.

Furthermore, the STRING score, which reflects the confidence of interaction based on aggregated experimental data and inferences from protein homology, exhibits a clear positive correlation with the predictions, especially when higher scores (red) were observed. Conversely, lower scores (blue) are associated with negative predictions. Detailed analysis of SHAP value and STRING score (Figure 2D) shows that the threshold for the feature contribution is in the medium confidence range with the STRING score of 350. Although the number of positive and negative instances for the GO match feature was lower than that for other features (Figure 2C), it still strongly influenced the prediction of true interactions. Collectively, these results demonstrate that incorporating robust large-scale biological features can enable the construction of prediction models capable of effectively evaluating mediocre hits and estimating the probability of true interactions. Overall, use of this approach resulted in an improved interactome which incorporated the predictions (Supplementary Figure S2).

### Structural mapping

#### Ribosome

To validate the reliability of our crosslinking data, we performed structural mapping of crosslinked residue pairs onto the experimentally determined 3D structures (Figure 3A and 3B; Supplemental Movie 1 and Supplemental Movie 2). Specifically, we utilized the cryo-EM structure of the 60S ribosomes (PDB: 5XXB) and 40S ribosomes (PDB: 5XXU), which had been previously resolved to near-atomic resolutions (37). For the two ribosome complexes, we mapped and measured the Cα-Cα distances of unique residue-residue inter- and intra-links, with a false discovery rate of less than 5%. Our analysis revealed that the median distances for inter-link was 21.8 Å (n=67) and intra-link is 16.5 Å (n=418). In contrast, the median distance for all possible KSTY residue pairs (the amino acids that could be linked by the crosslinker) was 111.0 Å (n=2324504), thereby demonstrating the validity of crosslinked pairs (Figure 3C).

**FIGURE 3:**
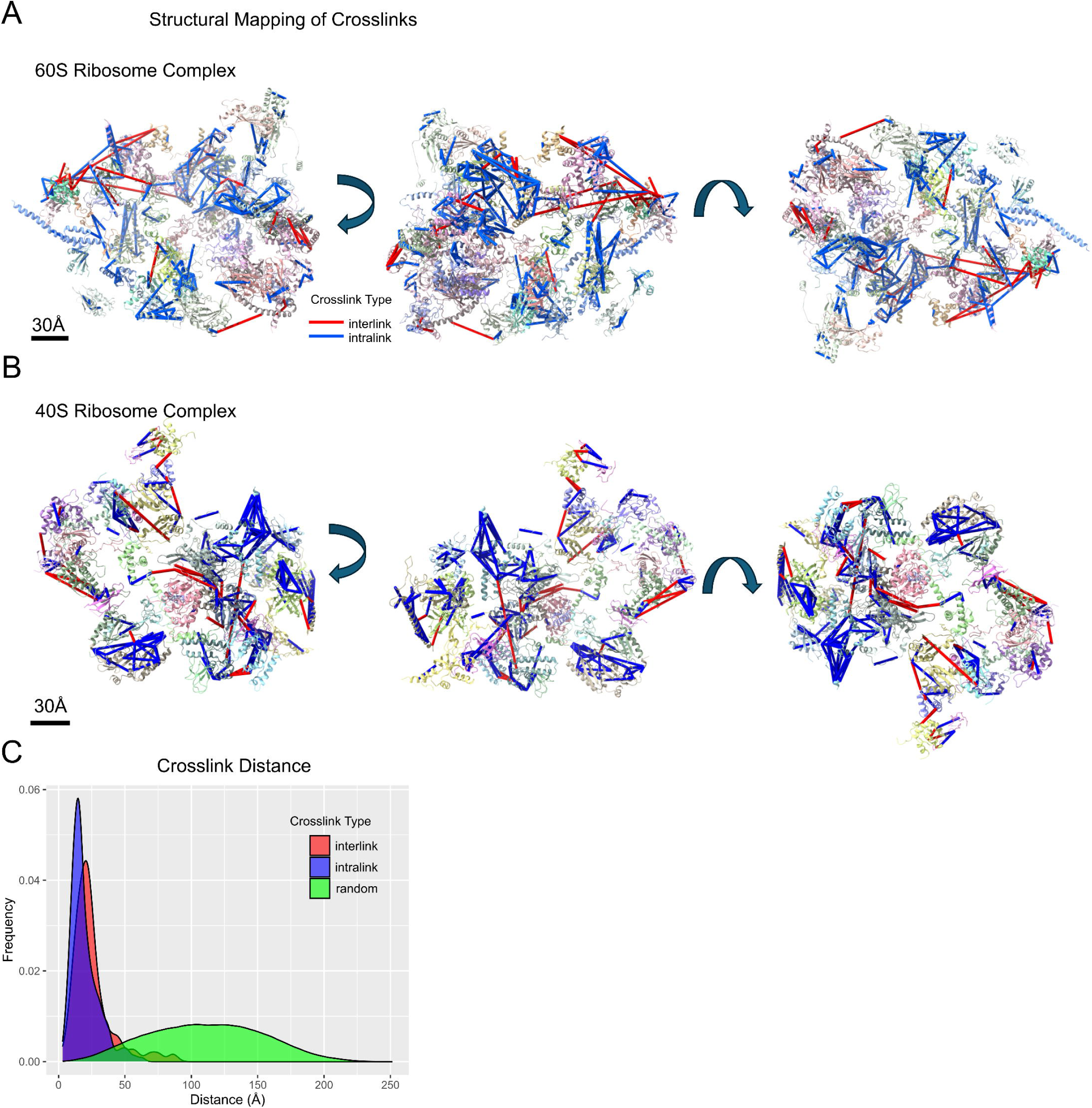
Structural Mapping of Ribosome Crosslinks. Interlinks (red) and intralinks (blue) were mapped onto previously determined cryo-EM structures of the 60S ribosome subunit complex **(A)** and the 40S complex **(B),** with three rotations shown for each structure. **(C)** A frequency histogram depicting the interlinks (red, M = 21.8 Å, n = 67), intralinks (blue, M = 16.5 Å, n = 418), and random KSTY-KSTY pairs (green, M = 111.0 Å, n = 2,324,504). This highlights the specificity of crosslinks relative to random pairs.

Due to weak local density and its transient association with the ribosome, the cryo-EM reconstruction did not resolve signaling scaffold protein RACK1 (TGME49_216880, annotated as POC1 centriolar protein in ToxoDB) within the 40S subunit. In contrast, our crosslinking data (Figure 4A) reveals that RACK1 interacts with RPS17, demonstrating that XL-MS can detect transient associations. Additionally, various ribosomal proteins (RPS20, RPS21, RPS29, PRSA, and RACK1) that were mischaracterized with HyperLOPIT data as cytosolic, chromatin, or unassigned are directly connected to the networks. This finding underscores the ability of XL-MS to complement and refine data obtained from other mainstream approaches. This data also demonstrates that XL-MS not only identified known protein interactions, but also found potential novel interactions. For example, the HEAT-repeat containing protein (TGME49_231600), which has homology to importin β, is crosslinked to ribosomal protein RPS23. This putative importin β might transfer nascent cytosolic RPS23 into the nucleus to be assembled with ribosomal RNA at the nucleolus as described in other organisms (38).

**FIGURE 4:**
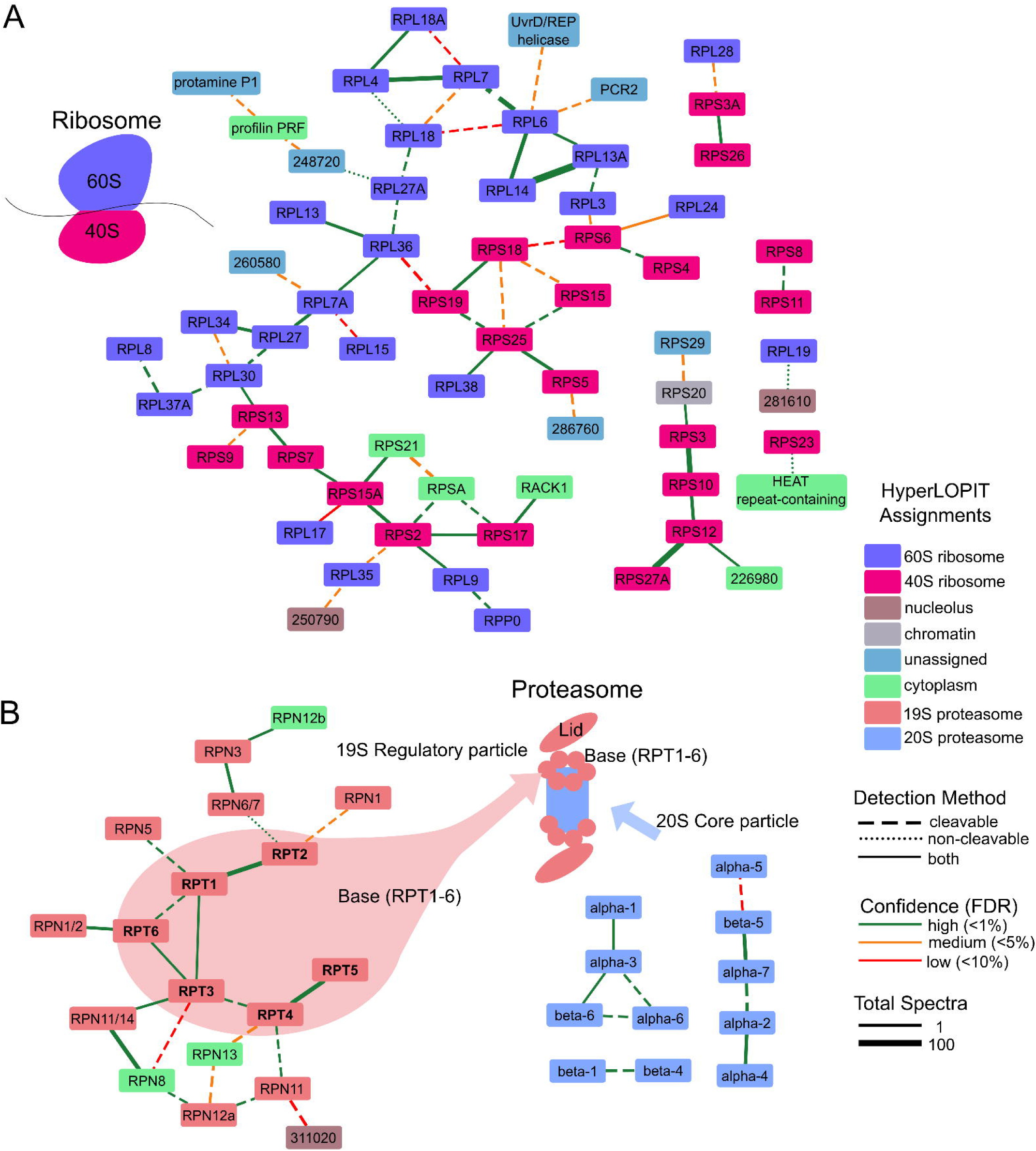
Ribosome and Proteosome Interactomes. The interactome for **(A)** ribosome-associated proteins and **(B)** proteasome proteins, with nodes color-coded by hyperLOPIT assignments. The network edges are color-coded based on confidence, with line styles indicating detection methods and thickness representing the total number of spectra for each interaction. These networks also integrate salvaged interactions identified through machine learning.

#### Proteasome complex

The proteasome consists of two major complexes: the 20S core particles, which function as the proteolytic shredder, and 19S regulatory particles. Our observations (Figure 4B) reveal a distinct segregation between these two assemblies. The 19S regulatory particle is further organized into two subcomplexes, the Lid and the Base. The Base comprises six AAA+ ATPases (Rpt1 through Rpt6; Figure 4B, Bold) that form a ring in a defined sequential order (39). Our crosslinking data not only confirms the ring structure but also corroborates the specific sequence of the ATPase subunits. Consistent with the ribosome findings, our XL-MS data revealed that three proteasomal proteins—RPN8, RPN12b, and RPN13—which were misclassified as cytosolic based on HyperLOPIT, are directly associated with the 20S core particle. These findings clarify the distinct structural organization of proteasome assemblies and reveal new functional associations, thereby enhancing our understanding of proteasomal regulation and protein degradation.

#### Homo-dimer detection

When two crosslinked peptide sequences overlap, it indicates that two identical protein molecules are in close proximity, suggesting the formation of homo-dimers. A thorough analysis of the peptide sequences identified 101 proteins that exhibited homo-dimer crosslinks of which 16 had 3 or more links (Table 4). Although direct evidence of homo-dimerization in *T. gondii* is scarce, many homologs in other organisms are known to form homodimers, such as peroxiredoxin 1 (40), phosphoenolpyruvate (PEP) carboxykinase I (41), coronin (42), fructose-1,6-bisphosphate aldolase (43), all of which are highly represented in the data set. These findings strongly support the validity of dimerization detection using cross link mass spectrometry data.

Interestingly, several GRA proteins were identified among the homodimer candidates, including GRA7, GRA2, GRA3, GRA32, GRA8, and GRA6. Specifically, GRA7 exhibited five unique residue pairs for homo-dimerization. Mapping the crosslinks revealed potential hotspots for dimerization (Figure 5A). Notably, GRA7 is known to form complexes with ROP18/ROP5 and to alter the oligomerization of Irga6 (44, 45). Similarly, GRA3 demonstrated three unique homodimerization residue pairs and is previously recognized for its capacity to dimerize, interact with the host Golgi apparatus, and facilitate membrane scavenging (46). These findings suggest that GRA protein dimerization may play a critical role in immune evasion and the appropriation of host cellular functions.

**FIGURE 5:**
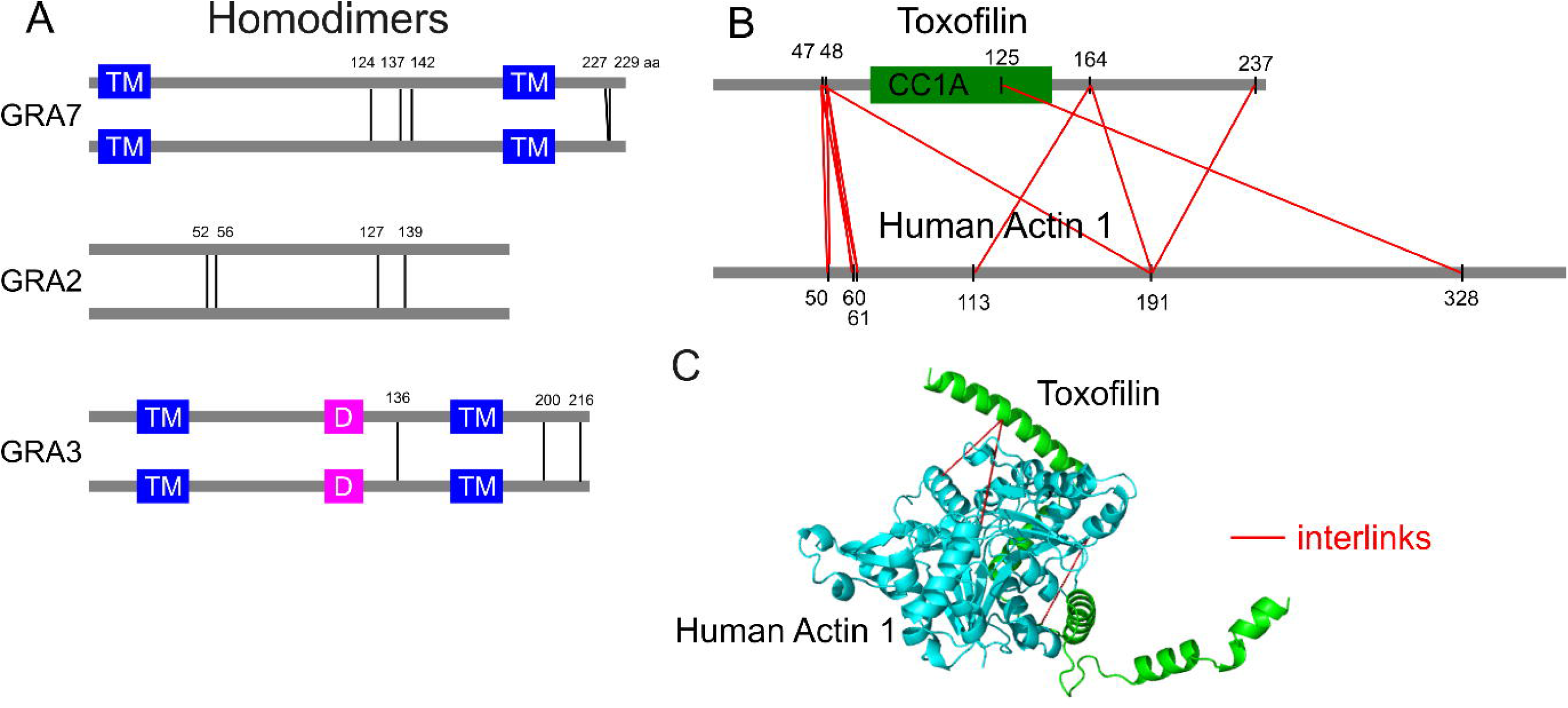
Protein Interactions. **(A)** Representative homodimers determined by crosslinking data are shown with protein amino acid positions and domain lengths drawn to scale (TM: transmembrane domain, D: dimerization domain). The lines indicate the residues involved in dimerization. **(B)** This panel depicts a representative human–parasite interaction at the residue-to-residue level, highlighting the CC1A domain in toxofilin, which is required for actin binding. **(C)** Structural mapping of toxofilin–actin crosslinks is presented on a 3D model based on homology modeling of a cryptographically determined structure (PDB: 2Q97) that accounts for the SNP present in toxofilin.

#### Host-parasite interactions

To evaluate the potential crosslinking events between *T. gondii* and human proteins, we re-analyzed our dataset using a combined database containing protein sequences from both organisms. Since the samples were prepared from purified parasite cytosolic lysates, which were intentionally depleted of host-derived material, very few human–parasite PPIs were identified. The most prominent pair observed was the interaction between toxofilin (TGME49_214080) and actin with eleven unique residue-to-residue crosslinks (Figure 5B). Toxofilin is known to bind to globular actin via its CC1A domain (47). Although several peptides were mapped to both human and parasite actin, other peptides were unique to the human sequence demonstrating the potential for dissecting human-parasite interactions. Three crosslinks were mapped onto the structure, which was homology modeled using the portion of toxofilin crystallized with human actin 1 (PDB: 2Q97), indicating that these crosslinks are consistent with the previously characterized host-pathogen complex (Figure 5C).

#### GRA Network

Unexpectedly, our study identified the dense granule protein network as a prominent subnetwork. This finding is noteworthy because the sample preparation was designed to process only cytosolic proteins. Considering that the majority of dense granule proteins are secreted into parasitophorous vacuoles, their presence in our samples likely occurs in small amounts, predominantly within dense granule membranous vesicles that “contaminated” the cytosolic preparation.

Figure 6A presents the selected interactome of known GRAs. Previous studies have established that GRA proteins within dense granules form oligomeric complexes exceeding 1 MDa throughout the secretory pathway (48). These pre-assembled oligomeric GRA complexes are thought to hide the hydrophobic domains, thus maintaining solubility in the dense granule matrix. Previous co-immunoprecipitation experiments (48) have demonstrated that GRA2 associates with GRA3, GRA6, and GRA7 (Figure 6B). Our crosslinking data further distinguishes between direct and indirect binding interactions. Notably, the interaction between GRA2 and GRA3 occurs indirectly through GRA7 (Figure 6A), offering intricate insights into complex assembly that were previously inaccessible through conventional assays. A more recent study examining the co-elution of native complexes in ToxoNET (6) produced a similar set of GRA proteins as potential complex components. In particular, ToxoNET Cluster #26 comprises GRA1, GRA2, GRA3, GRA4, GRA6, GRA7 (Figure 6B), and additional microneme proteins. Our data unambiguously delineates the specific binding relationships among these proteins.

**FIGURE 6:**
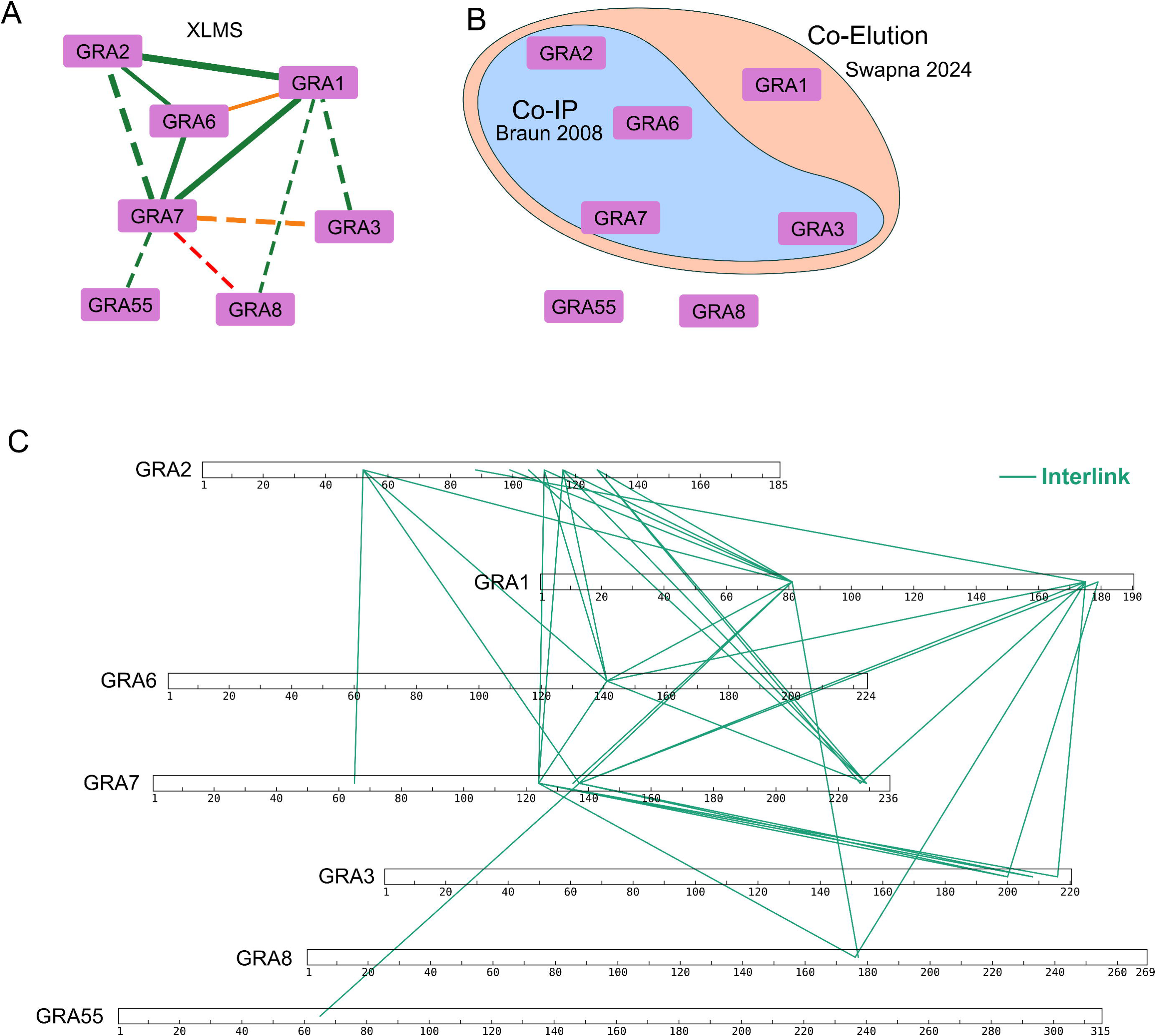
Dense granule (GRA) Protein Interactions Identified by Cross Linked Mass Spectrometry. **(A)** GRA interactome based on crosslinking data **(B)** GRA interactome derived from previous studies using co-immunoprecipitation with GRA2 (Braun et al. 2009) (48) and large-scale co-elution study (ToxoNET, Swapna et al. 2024) (6). **(C)** The GRA interactome presented at the residue-to-residue level, drawn to scale, with green lines representing interlinks.

To enhance the resolution of the complex, individual residue-to-residue interactions were mapped for each protein (Figure 6C). GRA2, for instance, is characterized by a central domain (88–127) that serves as a hotspot, engaging 12 unique interactions with GRA1, GRA6, and GRA7. Although GRA2 was previously considered the hub of this complex, our interactome analysis indicates that GRA7 exhibits a greater number of connections, thereby possibly functioning as the central hub. These findings underscore the utility of XL-MS in elucidating complex, large-scale protein-protein interactions.

## Discussion

The current study demonstrates that XL-MS can effectively map the complex network of protein–protein interactions within the cytosolic fraction of *T. gondii*. By integrating both MS-cleavable and non-cleavable spectra, we identified thousands of residue–residue contacts that validated known assemblies, such as ribosomal and proteasomal complexes, and uncovered novel interactions, particularly among dense granule proteins. Structural mapping further confirmed the spatial feasibility of the observed crosslinks, while a machine learning model leveraging diverse biological features salvaged interactions from mediocre-score hits by distinguishing true associations from potential false positives. Overall, these results reinforce the robustness of XL-MS as a tool for interactome analysis and provide fresh insights into the molecular architecture and functional dynamics of *T. gondii*.

A limitation of our experimental design is that we performed no additional fractionation beyond cytosol extraction. This restriction limits the majority of detection to highly abundant proteins— often estimated to represent roughly the top 20% of the proteome—due to their high dynamic range. Incorporating further subcellular fractionation techniques, such as hyperLOPIT, or affinity purification methods like immunoprecipitation or BioID, would significantly enhance proteomic coverage by distributing and resolving these abundant proteins, allowing better detection of proteins of lower abundance.

Additionally, we applied machine learning to develop a model that distinguishes potential true hits among mediocre-score hits. However, comprehensive biological features are available for only a limited number of organisms. Neglected eukaryotic parasites, such as Microsporidia, may not exhibit sufficient biological features for effective machine learning application. Nonetheless, as more systematic data become available, the use of predictive models could prove valuable for a broader range of organisms

The development of this crosslinking mass spectrometric protocol will allow additional investigations of the various protein interactomes in *T. gondii*. For example, the capacity of this parasite to persist within the host over extended periods is a key factor in its transmission and ability to cause recrudescent disease in immune compromised hosts. The cyst wall, which serves as the interface between the cyst and the host, plays a critical role in the pathogenesis of chronic toxoplasmosis. Until recently, it was impossible to maintain the bradyzoite culture for long periods of time in tissue culture and develop a structurally distinct thick cyst wall layer present in the cysts from mouse brains. The use of myotubes as a host enabled the development of large mature cysts (28 days post infection) with thick wall which developed broad tolerance to various antiparasitic drugs as well as acid/pepsin treatment (49, 50). The ability to generate using *in vitro* systems mature cysts, instead of isolating mouse derived brain cysts, will facilitate the use of XL-MS to study the composition and protein interactome of the *T. gondii* cyst wall.

While our crosslinking studies have identified numerous potential interactions, including those between parasite and human proteins, these findings will require further experimental validation. Future investigations employing complementary techniques, such as co-immunoprecipitation, yeast two-hybrid assays, or Förster resonance energy transfer (FRET), should be undertaken to confirm the predicted interactions and further elucidate their functional significance.

## Table Legends

**TABLE 1:** The number of crosslinks identified from *T. gondii* cytosolic proteins, as determined by both cleavable and non-cleavable search approaches applied to the same dataset. The combined use of these search strategies resulted in a 54–92% increase in the detection of unique residue-to-residue interactions. Abbreviations: CMS, crosslink spectra match; PPI, protein–protein interactions.

**TABLE 2:** Gene ontology (GO) term analysis of the crosslinked proteins was performed, with terms ranked by p-value, revealing a strong representation of typical cytosolic proteins. This analysis utilized ToxoDB version 67, and p-values were calculated using Fisher’s exact test.

**TABLE 3:** Summary of the percentage agreement between crosslinked pairs and multiple independent biological datasets. Percentages are stratified by crosslink confidence level, highlighting that crosslinks with higher confidence exhibit greater agreement. Crosslinked pairs with missing data were excluded from these calculations.

**TABLE 4:** A list of homodimers detected from the crosslinking data, including only proteins with homo-dimer crosslinking peptides containing at least three unique residue-to-residue interactions (#RRI). False detections due to tandem repeats have been removed.

**Supplemental Figure S1. SEC Fractions**

The number of crosslinks (CSM) and non-crosslinks (PSM) identified in the each fractions after SEC fractionation. The fraction from 1.18 ml to 1.23 ml had highest ratio (46.6) of corsslink/non-crosslink while the fraction from 1.23 ml to 1.28 ml had highest number (535) of crosslinks. The trace is the UV absorbance at 214 nm representing the amount of peptides.

**Supplementary Figure S2. *T. gondii* protein interactome.**

The complete interactome of *T. gondii* cytosolic proteins, integrated with machine learning predictions. Prediction thresholds were stratified based on confidence levels, with high-confidence predictions set at >0.2, medium at >0.5, and low at >0.6. The image is provided in high resolution, ensuring that all node labels are clearly legible.

**Supplemental Movie 1 and 2**

Movies illustrating crosslinks mapped onto cryo-EM structures of the 60S large ribosomal subunit (PDB: 5XXB, Supplemental Movie 1) and the 40S small ribosomal subunit (PDB: 5XXU, Supplemental Movie 2). Interlinks are represented by red lines, whereas intralinks are indicated by blue lines.

**Supplemental File 1:**

This file includes the various scripts utilized in this study. Crosslink processing and visualization were conducted using R with the tidyverse package, while machine learning analyses were executed using Python with LightGBM.

## Acknowledgements

We thank ToxoDB and STRING for providing essential resources for our bioinformatic analyses. We are also grateful to Fan Liu and Ying Zhu from the Leibniz-Forschungsinstitut für Molekulare Pharmakologie (Berlin, Germany) for their valuable discussions on crosslinker quality. We express our gratitude to Liang Zhao from the City University of New York (CUNY) for his guidance on machine learning and to Lamisha Shah (CUNY), Andrews Afrifa (CUNY), Narito Tsumuraya (Fair Lawn High School, NJ), and Kenju Tomita (Irvington High School, NY) for their significant contributions during the initial stages of our machine learning work. This project was supported by NIH grant RO1 AI134753 (LMW).

