## Supplementary figures and images for "Mapping a *Toxoplasma gondii* interactome by crosslinking mass spectrometry and machine learning"

### Figure S1

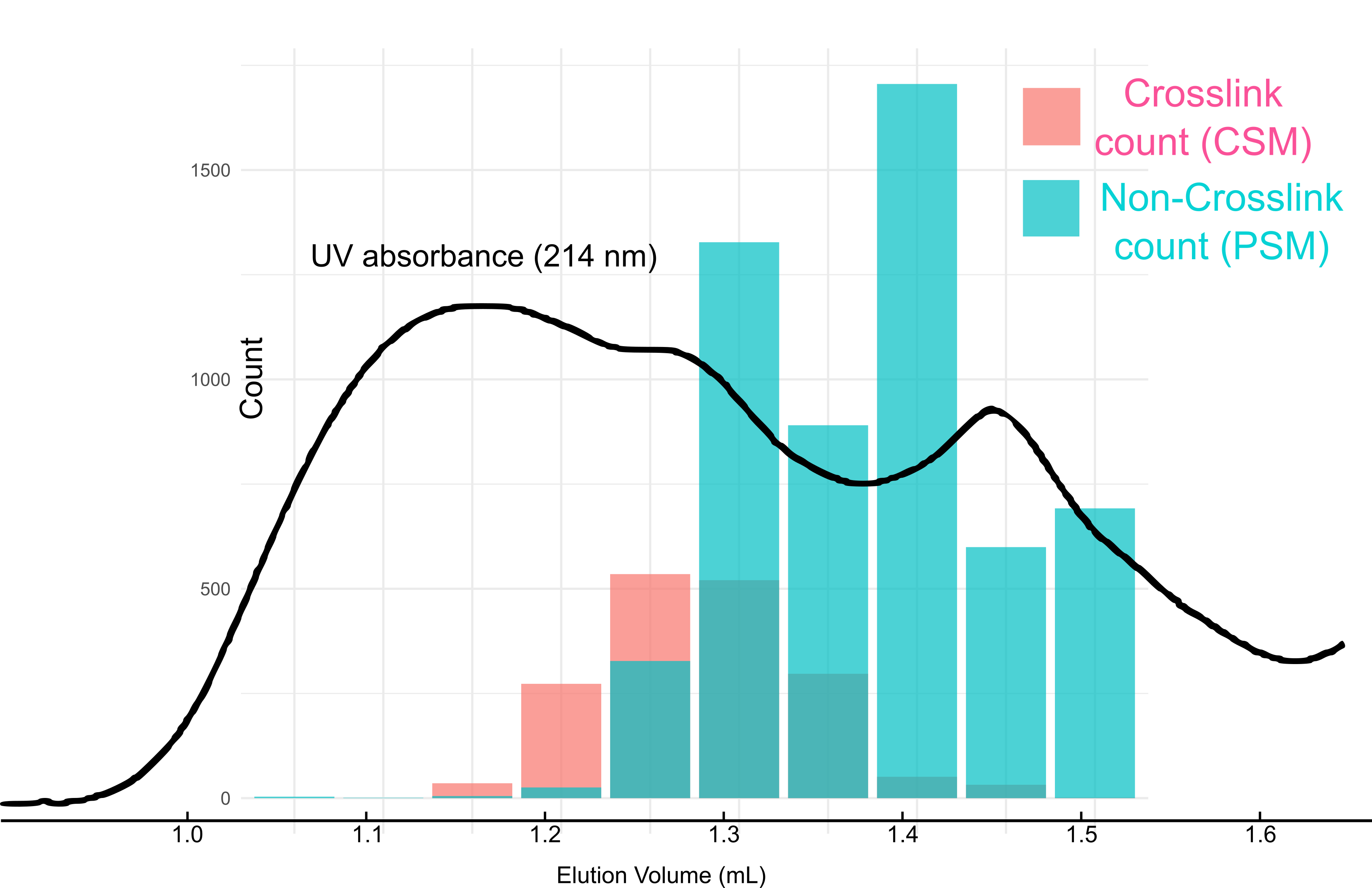

### Figure S2

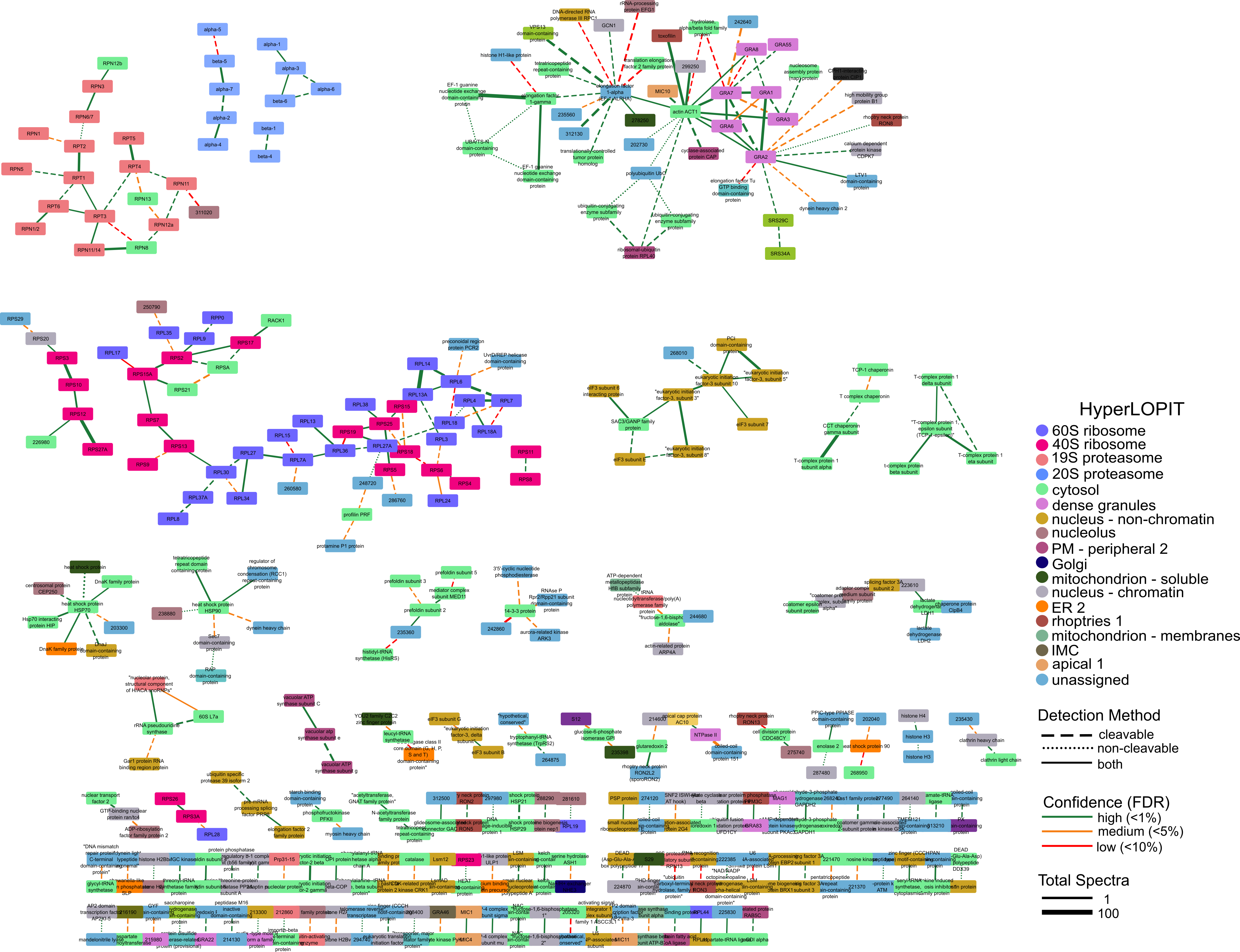
