## Supplementary File 1 for "Mapping a *Toxoplasma gondii* interactome by crosslinking mass spectrometry and machine learning"

---
title: "XL-MS processing and analysis using R"
output: html_notebook
---


```{r}
library(tidyverse)
```


#Load the clean full XLMS dataframe 
```{r}
#read the "crosslinks = residue pairs" results from the Annika data
rawdf <- read.table("TachyAll_XL_publication_20240916_Crosslinks.txt", header = TRUE, sep = "\t")

#removes suffix on protein names for Cytoscape
rawdf <- rawdf %>%
  mutate(across(where(is.character), ~str_replace_all(., "-t26_1-p1", ""))) %>%
  #remove unnecessary columns
  select(-c(Crosslinks., Checked, Crosslinker, Number.of.Proteins,))

#make the descriptions easy to read
# Define a function to clean specified columns with specific patterns, prefix removal, and custom string removals
clean_gene_product <- function(data, columns) {
  # Apply multiple transformations within mutate across the specified columns
  data %>% 
    mutate(across(all_of(columns), ~if_else(
      str_starts(., "transcript"),  # Check if starts with "transcript"
      str_split(., "\\|", simplify = TRUE)[, 4],  # Split and select the 4th element if condition is met
      .  # Keep the original value if condition is not met
    ))) %>%
    mutate(across(all_of(columns), ~str_remove(., "^\\s*gene_product="))) %>%  # Remove the "gene_product=" prefix
    mutate(across(all_of(columns), ~str_remove(., "dense granule protein "))) %>%  # Remove "dense granule protein " prefix
    mutate(across(all_of(columns), ~str_remove(., "ribosomal protein "))) %>%  # Remove "ribosomal protein " prefix
    mutate(across(all_of(columns), ~str_remove(., "SAG-related sequence "))) %>%  # Remove "SAG-related sequence
    mutate(across(all_of(columns), ~str_remove(., "heat shock protein ")))  # Remove "heat shock protein " prefix
}

# apply that funciton to df
rawdf <- clean_gene_product(rawdf, c("Protein.Descriptions.A", "Protein.Descriptions.B"))

```

#Since there are multiple genes matching to one peptide. Choose the first Toxo gene if it exist. 
```{r}
rawdf <- rawdf %>%
  separate_rows(Accession.A, sep = ";") %>% # Split gene names into separate rows
  group_by(Crosslinks.Peptide.Group.ID) %>% # Group by the original identifier
  arrange(!startsWith(Accession.A, "TGME49_")) %>% # genes starting with "TGME49_" come first
  slice_head(n = 1) %>%  # Select the first row of each group
  ungroup() %>%
  separate_rows(Accession.B, sep = ";") %>% #Perform the same with Accession.B
  group_by(Crosslinks.Peptide.Group.ID) %>%
  arrange(!startsWith(Accession.B, "TGME49_")) %>%
  slice_head(n = 1) %>%
  ungroup() %>%
  mutate(                                     #remove space from the end
    Accession.A = str_trim(Accession.A),
    Accession.B = str_trim(Accession.B),
    # Truncate everything after (and including) the first period "."
    Accession.A = str_remove(Accession.A, "\\..*"),
    Accession.B = str_remove(Accession.B, "\\..*")
  )

```

#Calculate Best.CSM.Score of FDR (corrected FDR calculation formula)
#Check the FDR at residue-residue level and score
%FDR 1, 5, 10, 30, 50
235 183 161 126 108 Tachy all 2024-09-16
284 213 191 118   cyst wall all
270 208 183 147 130 Toxo+Human all 2024-09-16
```{r}

df <- rawdf
# Initialize result data frame
results <- tibble(Threshold = integer(), Count_D = integer(), Count_T = integer())
# Maximum value in the Best.CSM.Score column
max_score <- max(df$Best.CSM.Score)
# Loop through thresholds
for (threshold in 1:max_score) {
  # Count occurrences of D
  count_D <- df %>%
    filter(Best.CSM.Score > threshold, Decoy == "True") %>%
    summarise(count = n()) %>%
    pull(count)
  # Count occurrences of T
  count_T <- df %>%
    filter(Best.CSM.Score > threshold, Decoy == "False") %>%
    summarise(count = n()) %>%
    pull(count)
  # Add the threshold and counts to the results data frame
  results <- results %>%
    add_row(Threshold = threshold, Count_D = ifelse(length(count_D) == 0, 0, count_D), Count_T = ifelse(length(count_T) == 0, 0, count_T))
}
results <- results %>%
  mutate(Percentage_D = ifelse((Count_D + Count_T) == 0, 0, 100 * Count_D / Count_T))

# Function to find nearest value (1, 5, 10, 50 %) and return corresponding Threshold scores
find_nearest_threshold <- function(target) {
  results %>%
    # Calculate the absolute difference from the target
    mutate(difference = abs(Percentage_D - target)) %>%
    # Arrange by this difference to bring the closest to the top
    arrange(difference) %>%
    # Slice the top row to get the closest value
    slice_head(n = 1) %>%
    # Select only the Threshold column
    pull(Threshold)
}

# Find nearest for each of the specific numbers
targets <- c(1, 5, 10, 30, 50)
FDRdf <- sapply(targets, find_nearest_threshold)
print(FDRdf)
```

#relabel the Confidence levels with Confidence.Q, make PPI and RRI
```{r}
thresholds <- FDRdf
# get rid of decoy
df <- rawdf %>% filter(Decoy == "False") %>%
# Modify 'Confidence' based on new conditions
  arrange(desc(Best.CSM.Score)) %>%
  mutate(Confidence = case_when(
    Best.CSM.Score > thresholds[3] ~ Confidence, # Keep original label
    Best.CSM.Score > thresholds[4] ~ "Purgatory", # Change to Purgatory
    Best.CSM.Score <= thresholds[4] ~ "Hell", # Change to Hell
    TRUE ~ Confidence # Default to keep original in all other cases
  ))
#add quantitative confidence
df <- df %>%
  mutate(Confidence.Q = case_when(
    Confidence == "High"      ~ 5, #FDR <1%
    Confidence == "Medium"    ~ 4, #FDR <5%
    Confidence == "Low"       ~ 3, #FDR <10%
    Confidence == "Purgatory" ~ 2, #FDR <30%
    Confidence == "Hell"      ~ 1, #FDR >30%
    TRUE                      ~ NA_integer_  # This line handles any unexpected values
  ))  %>%
  mutate(MS = "cleaved")%>% #add MS method as cleaved
  
  # Create Residue to Residue Interaction (RRI)
  rowwise() %>%  # Apply operations row-wise
  mutate(
  RRA = paste(Accession.A, In.protein.A, sep = ":"),
  RRB = paste(Accession.B, In.protein.B, sep = ":")
  ) %>%
  ungroup() %>%
  rowwise() %>%
  mutate(RRI = paste(sort(c(RRA, RRB)), collapse = "-"))%>% #sort alphanumerical then combine to make RRI
  ungroup() %>% # Remove row-wise grouping
  select(-c( RRA, RRB))%>% #remove non-relevent columns
  
  #consolidate observations with peptides of different length with same RRI into one
  group_by(RRI) %>%
  mutate(Number.of.CSMs = sum(Number.of.CSMs)) %>%   # sum the number of CSM.
  slice_max(order_by = Best.CSM.Score, n = 1, with_ties = FALSE) %>% # keep the highest CSM score observation
  ungroup() %>%

  #Create protein protein interaction (PPI)
  rowwise() %>%  # Apply operations row-wise
  mutate(
    # Combine accessions into a vector, sort them, and paste with "-"
    PPI = (paste(sort(c(Accession.A, Accession.B)), collapse = "-")
  )) %>%
  ungroup()
```


###get the summary of the counts in the confidence intervals

```{r}
#count each interval and print out the summary
countdf <- df %>% filter(Decoy == "False") %>% count(Confidence.Q) %>% arrange(desc(Confidence.Q)) 

combined_df <- cbind(targets, FDRdf, countdf) %>% 
  rename(
        `FDR_%` = targets,
        cutoff_score = FDRdf,
        interval_count = n
          ) %>% # Add a new column with the cumulative sum of 'interval_count'
  mutate(cumulative_count = cumsum(interval_count)) %>%
  mutate(cumulative_false_count = `FDR_%` * cumulative_count / 100) %>%
  # Convert cumulative count to interval count
  mutate(interval_false_count = cumulative_false_count - lag(cumulative_false_count, default = cumulative_false_count[1]))%>%
  mutate(`interval_FDR_%` = interval_false_count / interval_count * 100) %>%
  mutate(`interval_TDR_%` = 100 - `interval_FDR_%`)
# Print the long format dataframe
print(combined_df)
```

### Coutn the number of cross links at FDR 1 and 5%
```{r}
# Create a new dataframe with the required counts
counts_df <- df %>%
  # Step 1: Filter out rows where Decoy is "False"
  filter(Decoy == "False") %>%
  
  # Step 2: Group by Crosslink.Type
  group_by(Crosslink.Type) %>%
  
  # Step 3: Summarise counts for High and High & Medium confidence
  summarise(
    High_Confidence = sum(Confidence == "High"),
    High_Medium_Confidence = sum(Confidence %in% c("High", "Medium")),
    .groups = "drop"  # Ungroup after summarising
  ) %>%
  
  # Step 4: Add a total row across all Crosslink.Type
  bind_rows(
    summarise(
      .,
      Crosslink.Type = "Total",
      High_Confidence = sum(High_Confidence),
      High_Medium_Confidence = sum(High_Medium_Confidence)
    )
  )

# View the resulting counts dataframe
print(counts_df)

```

#visualize the decoy/true in cumulative counts
```{r}
# Reshape the data to long format for plotting
long_results <- results %>%
  pivot_longer(cols = c(Count_D, Count_T), names_to = "Decoy", values_to = "Count")

# Create the plot
ggplot(long_results, aes(x = Threshold, y = Count, color = Decoy)) +
  geom_line() +
  labs(title = "Counts of Decoy and T at Different Thresholds",
       x = "best CSM score",
       y = "Count",
       color = "Decoy Type") +
  theme_minimal()+
   # Add vertical lines at the values in FDRdf
  geom_vline(xintercept = FDRdf, linetype = "dashed", color = "green")+
  # Limit the Y-axis to 20000
  ylim(0, 20000)+
  #limit the X-axis to 700
  xlim(0, 700)
```

### insted of frequency (geom_density), use histogram (counts) to reflect the hight of the real hits and decoy hits

```{r}
# Create a plot showing the count of real and decoy hits
gg<- ggplot(rawdf, aes(x = Best.CSM.Score)) +
  # Plot the stacked histogram of True/False in Decoy (non-stacked!)
  geom_histogram(aes(fill = Decoy), binwidth = 5, position = "identity", alpha = 0.5) +
  labs(title = "Count of Real/Decoy hits vs Best CSM Score",
       x = "Best CSM Score",
       y = "Counts",
       fill = "Decoy") +
  theme_minimal()+
   # Add vertical lines at the values in FDRdf
  geom_vline(xintercept = FDRdf[1:3], linetype = "dashed", color = "black")+
  # Limit the Y-axis to 20000
  ylim(0, 2000)+
  #limit the X-axis to 700
   xlim(90, 600)+
  # Define custom colors: red for Decoy and light blue for Real
  scale_fill_manual(values = c("True" = "orange", "False" = "blue"),
                    labels = c("True" = "Decoy","False" = "Real"))
print(gg)
# Export the plot as SVG
ggsave("histogram_decoy_vs_true_plot.svg", plot = gg, width = 5, height = 3, device = "svg")
```

#keep the uncondensed (has multiple residue-to-residue)
```{r}
RRI_cleaved_full <- df
#uncondensed_df <- df
#Write the uncondensed data into a text file #Excel can't hold too many characters per cell. Use text insted.
write.table(RRI_cleaved_full, file = "TachyAll_XL_publication_20240916_uncondensed_woFeatures.txt", sep = "\t", row.names = FALSE, col.names = TRUE)

#revive uncondensed
#df <- read.table(file = "TachyAll_XL_publication_20240916_uncondensed_woFeatures.txt", sep = "\t", header = TRUE)
```

#Condense PPI network data for Machine Learning 
#calculate number of unique intra link for each protein
#remove decoy, remove intra, create PPI ID, count number of uniuqe residue-residue crosslinks, keep highest score, and collapse redundancy. Also add Total CSM numbers.
```{r}
#calculate number of unique intralinks for each protein at FDR <10% to use for feature
intracalc <- df %>%
    filter(Decoy == "False") %>%
    filter(Confidence.Q >= 1) %>% # include all #This is FDR of <10%
    filter(Crosslink.Type == "Intra") %>%
  group_by(Accession.A) %>%
  mutate(number_unique_intra = n()) %>%
  ungroup()%>%  # Optionally, remove the grouping information
  group_by(Accession.A) %>%
    # Select the row with the highest 'Best.CSM.Score' for each 'PPI'
    filter(Confidence.Q == max(Confidence.Q)) %>%
    ungroup() %>%
    distinct(Accession.A, .keep_all = TRUE)%>% #remove redundancy
    select(Accession.A, number_unique_intra) #select ID and interlink number

#collapse unique residue-residue to PPI
df <- df %>% 
    filter(Decoy == "False") %>%
    filter(Crosslink.Type == "Inter") %>%
  rowwise() %>%
    mutate(PPI = paste(sort(c(Accession.A, Accession.B)), collapse = "-")) %>%
    ungroup() %>%
  group_by(PPI) %>%
  mutate(Total.CSMs = sum(Number.of.CSMs),
         #crosslinks for XlinkCyNET format
        crosslinks_ab = paste(Accession.A, In.protein.A, Accession.B, In.protein.B, sep = "-", collapse = "#"),
        score_ab = paste(Best.CSM.Score, collapse = "#")
        )%>%
  ungroup()%>%  # Optionally, remove the grouping information
  group_by(PPI) %>%
    mutate(number_unique_RR = n()) %>% #count unique residue-residue pair
    # Select the row with the highest 'Best.CSM.Score' for each 'PPI'
    filter(Best.CSM.Score == max(Best.CSM.Score)) %>%
    ungroup() %>%
    distinct(PPI, .keep_all = TRUE)%>% #remove redundancy
    select(-Number.of.CSMs) #remove the number of CSM since it's not representing after collapse
#append unique intralinks (keep the smaller of two to represent the uncertinity of the peptide)
df <- df %>%
  left_join(intracalc, by = "Accession.A") %>%
  left_join(intracalc, by = c("Accession.B" = "Accession.A")) %>%
  # Replace NA with 0 and compute the minimum of 'number_unique_intra.x' and 'number_unique_intra.y'
  mutate(number_unique_intra.min = pmin(replace_na(number_unique_intra.x, 0), replace_na(number_unique_intra.y, 0), na.rm = TRUE)) %>%
  mutate(number_unique_intra.hm = 2 * (number_unique_intra.x * number_unique_intra.y)/(number_unique_intra.x + number_unique_intra.y)) %>%
  # Remove the old columns 'number_unique_intra.x' and 'number_unique_intra.y'
  select(-number_unique_intra.x, -number_unique_intra.y)
```

### coutn the number of PPI at FDR 1 and 5%
```{r}

# Create a new dataframe with the required counts
counts_df <- df %>% #decoy is already filterd out
  # Step 2: Group by Crosslink.Type
  group_by(Crosslink.Type) %>%
  
  # Step 3: Summarise counts for High and High & Medium confidence
  summarise(
    High_Confidence = sum(Confidence == "High"),
    High_Medium_Confidence = sum(Confidence %in% c("High", "Medium")),
    .groups = "drop"  # Ungroup after summarising
  ) %>%
  
  # Step 4: Add a total row across all Crosslink.Type
  bind_rows(
    summarise(
      .,
      Crosslink.Type = "Total",
      High_Confidence = sum(High_Confidence),
      High_Medium_Confidence = sum(High_Medium_Confidence)
    )
  )

# View the resulting counts dataframe
print(counts_df)
```

#Getting Features (skip if using the FeaturesDF from a backup file)
#Features from ToxoDB and ToxoNET
```{r}
featureDF <- read_excel("ToxoDB67_GO_TGME49.xlsx") %>%
  rename_with(~ gsub(" ", "_", .x))%>% #excel has space in the column name. replace with "_"
  rename_with(~ gsub("/", "_", .x)) #also remove "/" from the column name
  # Function to concatenate unique values of a vector
concat_unique <- function(x) {
  unique_x <- unique(x)
  str_c(unique_x[!is.na(unique_x)], collapse = ";")
}

featureDF <- featureDF %>%
  group_by(Gene_ID) %>%
  summarise(across(everything(), ~concat_unique(.x), .names = "{.col}_c"), .groups = 'drop')

# Function to remove redundancy within each cell and concatenate back
clean_values <- function(x) {
  # Ensure that x is a character vector
  x <- as.character(x)
  # Apply the cleaning process
  str_split(x, pattern = ";") %>% 
    map(~.x[.x != "N/A"]) %>%  # Remove "N/A" values
    map(~.x[.x != "GO:0016020"]) %>%  # Remove the specific element "GO:0016020"
    map(unique) %>%  # Remove duplicate values
    map_chr(~str_c(.x, collapse = ";"))
}

# Apply the function across all columns of the dataframe
featureDF <- featureDF %>%
  mutate(across(everything(), clean_values))

#combine CRISPR data to featureDF
CRISPRdf <- read_excel("ToxoDB67_CRISPR.xlsx")%>%
  rename_with(~ gsub(" ", "_", .x))%>% #replace space with "_"
  mutate(Gene_ID = str_replace_all(Gene_ID, "TGGT1_", "TGME49_"))
featureDF <- left_join(featureDF, CRISPRdf, by = "Gene_ID")

#load ToxoNET data and combine to featureDF
ToxoNETdf <- read_excel("ToxoNETclusters.xlsx")%>%
  separate_rows(Components, sep = ",")%>%
  select(Components, Cluster)%>%
  group_by(Components) %>% summarise(Cluster = paste(unique(Cluster), collapse = ";")) %>% 
  ungroup()%>%
  rename(Gene_ID = Components, ToxoNET = Cluster) #rename column names

#append ToxoNET feature to the main feature data frame
featureDF <- left_join(featureDF, ToxoNETdf, by = "Gene_ID")

featureDF <- featureDF %>% 
  select(-source_id_c, -Product_Description_c)%>%
  rename(RnaSeq = "Stage-specific_RNA-sequencing_in_Toxoplasma_gondii_-_24h_tachyzoites_-_unique_(Toxoplasma_gondii)_c")

#load hyperLOPIT data and add to featureDF
LOPITdf <- read_excel("ToxoDB67_LOPIT_TGME49.xlsx")%>%
  rename_with(~ gsub(" ", "_", .x))%>% #replace space with "_"
  select(Gene_ID, Location)%>%
  filter(Location != "outlier")%>% #remove outliers
  group_by(Gene_ID) %>% summarise(Location = paste(unique(Location), collapse = ";")) %>% ungroup()
featureDF <- left_join(featureDF, LOPITdf, by = "Gene_ID")
featureDF <- featureDF %>% mutate(across(everything(), ~na_if(.x, ""))) #add na to empty cells

#STRING!!!!!!!!!!!!!!!!!!!!!!!!!!!!!!!!!!!!!!!!!!!!!!!!!!!!!!!!!!!!!!!!!!!!!!!!!!!!!!!!!!!!!!!!!!!
###############perform only once to get all STRING names and searh with UniProt ID Mapping#########
# STRINGdf <- read.table("432359.protein.links.detailed.v12.0.txt", header = TRUE, sep = " ")
# STRINGnames <- STRINGdf %>%
#   # Extract the part after the "." in both columns
#   mutate(across(c(protein1, protein2), ~str_extract(., "(?<=\\.).+"))) %>%
#   # Transform the data to a long format
#   pivot_longer(cols = c(protein1, protein2), values_to = "STRINGnames") %>%
#   # Select only the STRINGnames column
#   select(STRINGnames) %>%
#   # Remove duplicate entries
#   distinct() %>%
#   # Optionally, you might want to remove NA values if any extraction failed
#   drop_na()
# # Combine all values in the "STRINGnames" column, separated by a space
# combined_string <- paste(STRINGnames$STRINGnames, collapse = " ")
# # Save the combined string to a text file -> use UniProt ID Mapping to get VEG names
# writeLines(combined_string, "STRINGnames.txt")

#read UniProt ID Mapped gene names for STRING
aliasdf <- read.table("STRINGnames_idmapping_2024_04_11.tsv", header = TRUE, sep = "\t") %>%
  mutate(To = str_extract(To, "(?<=:).*")) %>% # Modify the "To" column to keep only the part after the ":"
  mutate(To = str_replace_all(To, "TGVEG_", "TGME49_")) %>% #replace the VEG to ME49 gene name
  mutate(To = str_replace(To, "[A-Z]$", ""))%>% #remove the last alphabet in the gene names
  rename(UPname = From, MEname = To) #change name to Uniprot name and ME49 name

#read STRING network data and shape it for processing
STRINGdf <- read.table("432359.protein.links.detailed.v12.0.txt", header = TRUE, sep = " ")%>%
  mutate(across(c(protein1, protein2), ~str_extract(., "(?<=\\.).+")))%>% # keep the part after the "."
  left_join(aliasdf, by = c("protein1" = "UPname")) %>% #add TG name for protein1
  rename(Accession.A= MEname) %>%
  left_join(aliasdf, by = c("protein2" = "UPname")) %>% #add TG name for protein2
  rename(Accession.B= MEname)  %>%
  filter(!is.na(Accession.A) & !is.na(Accession.B)) %>% #remove na containing rows
  rowwise() %>%
    mutate(PPI = paste(sort(c(Accession.A, Accession.B)), collapse = "-")) %>% #join two proteins to make PPI ID
    ungroup() %>%   
  select(PPI, combined_score)%>% #just get the PPI and score
  distinct(PPI, .keep_all = TRUE)%>% #just keep the unique (remove 2x redundancy)
  rename(STRING.score = combined_score) #rename the column name score to reflect STRING
#load STRING data to Feature DF
write.table(STRINGdf, file = "ToxoDB67_AllSTRING.txt", sep = "\t", row.names = FALSE, col.names = TRUE) 

#Write the features into a text file
write.table(featureDF, file = "ToxoDB67_AllFeature_woSTRING.txt", sep = "\t", row.names = FALSE, col.names = TRUE)
```


##### pLink search processing ####
#perform pLink search with FDR at 10, 5, 1%
#pLink peptide-peptide chart has incovenitent shape to read and use for downstream. Use "Peptide_Order" as identifyer, separate summary from observation, find the max scores, calculate total PSM, and append to summary.
```{r}
# Define the column names (replace with meaningful names if available)
column_names <- c("Peptide_Order", "Protein", "Unique_Peptide_Number", "Number.of.CSMs", "Precursor_Mass", "Peptide", "Peptide_Mass",                   "Modifications", "Evalue", "Score", "Da", "ppm", "Alpha_Evalue", "Beta_Evalue")
#read the pLink full data (largest one)
pLinkrawdf <- read.table(
  "pLink_cross-linked_sitesFDR10.csv", 
  #"pLink_cross-linked_sitesFDR05.csv", 
  #"pLink_cross-linked_sitesFDR01.csv", 
                  col.names = column_names, 
                  header = FALSE, 
                  sep = ",", 
                  fill = TRUE, 
                  stringsAsFactors = FALSE,
                  na = c("", "NA"),
                  skip = 2) %>%
#removes suffix on protein names
  mutate(across(where(is.character), ~str_replace_all(., "-t26_1-p1", "")))%>%
  mutate(Protein = str_remove_all(Protein, "\\.P\\d{2,4}"))%>%
#convert probablity score to -log10
  mutate(Score = -log10(Score))%>%
#from the summary rows, add the Peptide_Order to the individual observations below
# Fill NA values in "Peptide_Order" with the previous non-NA value. This will be the key ID for two data frames
  mutate(Peptide_Order = str_replace_all(Peptide_Order, "[^0-9\\.]", "") %>% as.numeric())%>% #remove "SameSet" characters
  fill(Peptide_Order, .direction = "down")

#create summary data frame by "Protein" column is character
summarydf <- pLinkrawdf %>% filter(str_detect(Protein, "^[A-Z]"))%>%
                        select(-c(Precursor_Mass,Peptide,Peptide_Mass, Da, ppm, Alpha_Evalue, Beta_Evalue, Modifications, Peptide_Mass, Evalue, Score))
#Create observationdf: All other rows. keep entire observations with the highest Score per Peptide_Order
observationdf <- pLinkrawdf %>%
  filter(!str_detect(Protein, "^[A-Z]")) %>%                # Exclude rows where Protein starts with a capital letter
  group_by(Peptide_Order) %>%                               # Group by Peptide_Order
  slice_max(order_by = Score, n = 1, with_ties = FALSE) %>% # Keep the row with the highest Score per group
  ungroup() %>%
  select(c(Peptide_Order, Peptide, Score ))
#combine the best CSM score's observation to the summary
merged_summary <- summarydf %>% left_join(observationdf, by = "Peptide_Order") %>%
#split the Protein into Accession.A and B
  separate_rows(Protein, sep = "/")%>%
  filter(!is.na(Protein) & Protein != "")%>%
  separate(Protein, into = c("Accession.A", "Accession.B"), sep = "-", remove = FALSE)%>%
  # Separate Accession.A into Accession.A and In.protein.A
  separate(
    Accession.A,
    into = c("Accession.A", "In.protein.A"),
    sep = " ",                # Split at the first space
    extra = "merge",          # Merge any additional splits into the second column
    fill = "right"            # If there's no second part, fill with NA
  ) %>%
  # Clean In.protein.A by removing parentheses and any remaining whitespace
  mutate(
    In.protein.A = str_remove_all(In.protein.A, "[\\(\\)]")
  ) %>%
  # Repeat the separation for Accession.B
  separate(
    Accession.B,
    into = c("Accession.B", "In.protein.B"),
    sep = " ",                # Split at the first space
    extra = "merge",          # Merge any additional splits into the second column
    fill = "right"            # If there's no second part, fill with NA
  ) %>%
  # Clean In.protein.B by removing parentheses and any remaining whitespace
  mutate(
    In.protein.B = str_remove_all(In.protein.B, "[\\(\\)]")
  ) %>%
  # Check if they are Intra or Inter crosslinks
  mutate(Crosslink.Type = if_else(
    Accession.A == Accession.B,   # Condition
    "Intra",                       # Value if TRUE
    "Inter"                        # Value if FALSE
  )) %>%
  #Create PPI by %>%
  rowwise() %>%  # Apply operations row-wise
  mutate(
    # Combine accessions into a vector, sort them, and paste with "-"
    PPI = (paste(sort(c(Accession.A, Accession.B)), collapse = "-")
  )) %>%
  ungroup() %>%  # Remove row-wise grouping
  # remove the redundant PPI due to the non-unique peptide (one sequence hits multiple proteins)
  group_by(Peptide_Order) %>%                   # Group by Peptide_Order
  slice_min(order_by = PPI, n = 1, with_ties = FALSE) %>%  # Select the first aplhanumerical PPI per group
  ungroup() %>%
  #create peptide sequence "Sequence.A" position in peptide "Position.A" and B to be compatible with Annika 2.0
  separate(Peptide, into = c("Accession1", "Accession2"), sep = "-") %>%
  mutate(
    Sequence.A = str_extract(Accession1, "^[A-Za-z]+"),
    Position.A = as.integer(str_extract(Accession1, "[0-9]+")),
    Sequence.B = str_extract(Accession2, "^[A-Za-z]+"),
    Position.B = as.integer(str_extract(Accession2, "[0-9]+"))
    )%>%
   #convert pLink format of the peptide to Annika 2.0 format with the target residue in []
   mutate(
    Sequence.A = paste0(
      str_sub(Sequence.A, 1, Position.A - 1),
      "[", 
      str_sub(Sequence.A, Position.A, Position.A),
      "]",
      str_sub(Sequence.A, Position.A + 1)
    ),
      Sequence.B = paste0(
      str_sub(Sequence.B, 1, Position.B - 1),
      "[", 
      str_sub(Sequence.B, Position.B, Position.B),
      "]",
      str_sub(Sequence.B, Position.B + 1)
    )
  ) %>%
  # Create Residue to Residue Interaction (RRI)
  rowwise() %>%  # Apply operations row-wise
  mutate(
  RRA = paste(Accession.A, In.protein.A, sep = ":"),
  RRB = paste(Accession.B, In.protein.B, sep = ":")
  ) %>%
  ungroup() %>%
  rowwise() %>%
  mutate(RRI = paste(sort(c(RRA, RRB)), collapse = "-"))%>% #sort alphanumerical then combine to make RRI
  ungroup() %>% # Remove row-wise grouping
  select(-c(Accession1, Accession2, RRA, RRB)) #remove non-relevent columns
```
Perform above script for FDR 10, 5, 1
save the data frame. Combine them by checking with unique RRI and give them the Confidence.Q

```{r}
#RRI_noncleaved_FDR10 <- merged_summary
#RRI_noncleaved_FDR05 <- merged_summary
#RRI_noncleaved_FDR01 <- merged_summary
#merge PPI with FDR 1, 5, 10 %. before processing, create the PPI_condensed_FDR01, 05, 10
RRI_noncleaved_full <- RRI_noncleaved_FDR10 %>%
  mutate(Confidence.Q = 3) %>%
  mutate(Confidence.Q = if_else(RRI %in% RRI_noncleaved_FDR05$RRI, 4, Confidence.Q)) %>%
  mutate(Confidence.Q = if_else(RRI %in% RRI_noncleaved_FDR01$RRI, 5, Confidence.Q)) %>%
  mutate(MS = "noncleaved")
#export for merging with cleavable data
write.table(RRI_noncleaved_full, "TachyAll_XL_publication_20240916_RRI.noncleaved.csv", sep = "\t", row.names = FALSE, quote = FALSE)
```

#Calculate the number of RRI
```{r}
# Compute counts for Total
total_counts <- RRI_noncleaved_full %>%
  summarise(
    FDR10 = sum(Confidence.Q >= 3),
    FDR05 = sum(Confidence.Q >= 4),
    FDR01 = sum(Confidence.Q == 5)
  ) %>%
  mutate(Category = "Total")

# Compute counts for Intra
intra_counts <- RRI_noncleaved_full %>%
  filter(Crosslink.Type == "Intra") %>%
  summarise(
    FDR10 = sum(Confidence.Q >= 3),
    FDR05 = sum(Confidence.Q >= 4),
    FDR01 = sum(Confidence.Q == 5)
  ) %>%
  mutate(Category = "Intra")

# Compute counts for Inter
inter_counts <- RRI_noncleaved_full %>%
  filter(Crosslink.Type == "Inter") %>%
  summarise(
    FDR10 = sum(Confidence.Q >= 3),
    FDR05 = sum(Confidence.Q >= 4),
    FDR01 = sum(Confidence.Q == 5)
  ) %>%
  mutate(Category = "Inter")

# Combine all counts into one table
summary_table <- bind_rows(total_counts, intra_counts, inter_counts) %>%
  select(Category, FDR10, FDR05, FDR01)  # Arrange columns in desired order

# Print the table
print(summary_table)


```

#revive uncondensed RRI cleaved full data frame
```{r}
RRI_cleaved_full <- read.table(file = "TachyAll_XL_publication_20240916_uncondensed_woFeatures.txt", sep = "\t", header = TRUE)
```

#Merge the cleaved and noncleaved RRI together for Cytoscape and structural mapping
```{r}
#merge cleaved/noncleaved for cytoscape
RRI_cleaved_full <- RRI_cleaved_full %>% mutate(MS = "cleaved")
RRI_noncleaved_full <- RRI_noncleaved_full %>% mutate(MS = "noncleaved") %>%
  #convert pLink format of the peptide to Annika 2.0 format with the target residue
   mutate(
    Sequence.A = paste0(
      str_sub(Sequence.A, 1, Position.A - 1),
      "[", 
      str_sub(Sequence.A, Position.A, Position.A),
      "]",
      str_sub(Sequence.A, Position.A + 1)
    ),
      Sequence.B = paste0(
      str_sub(Sequence.B, 1, Position.B - 1),
      "[", 
      str_sub(Sequence.B, Position.B, Position.B),
      "]",
      str_sub(Sequence.B, Position.B + 1)
    )
  )

RRI_full <- full_join(RRI_cleaved_full, RRI_noncleaved_full, join_by(RRI))%>%
  mutate(
    MS = case_when( #MS is the MS detection methods
      !is.na(MS.x) & !is.na(MS.y) ~ "both",  # Both values exist
      !is.na(MS.x) & (is.na(MS.y)) ~ MS.x, # Only MS.x exists
      is.na(MS.x) & !is.na(MS.y) ~ MS.y,                   # Only MS.y exists
      TRUE ~ NA_character_))%>%
  #select larger of Confidence Q between cleaved and uncleaved
  mutate(Confidence.Q = pmax(Confidence.Q.x, Confidence.Q.y, na.rm = TRUE)) %>%
  #Add Number of CSMs
  mutate(Number.of.CSMs = rowSums(select(., Number.of.CSMs.x, Number.of.CSMs.y), na.rm = TRUE))%>%
  #get rid of non-relevant columns
  select(-c(Confidence, Confidence.Q.x, Confidence.Q.y, Crosslinks.Peptide.Group.ID, 
            Peptide_Order, Number.of.CSMs.x, Number.of.CSMs.y, MS.x, MS.y, 
            Protein.Descriptions.A, Protein.Descriptions.B)) %>%
  # Replace NA in all .x columns with corresponding .y columns
  mutate(across(ends_with(".x"), ~ coalesce(.x, get(sub("\\.x$", ".y", cur_column()))))) %>%
  # rename all columns that end with .x and remove the .x from the name 
  rename_with(
    .fn = ~ str_replace(., "\\.x$", ""),  # Function to remove '.x' at the end
    .cols = ends_with(".x")                # Select columns that end with '.x'
              ) %>%
  #select usable columns and clean up
  select(c(RRI, PPI, Accession.A, Accession.B, Confidence.Q, Number.of.CSMs, MS, Sequence.A, Position.A, Sequence.B, Position.B, In.protein.A, In.protein.B, Crosslink.Type, Best.CSM.Score))

#export for downstream analysis
write.table(RRI_full, "TachyAll_XL_publication_20240916_RRI.full.csv", sep = "\t", row.names = FALSE, quote = FALSE)
```

#####XMAS formatting for structural mapping################################
```{r}
#rearrange the columns for Xlinkx format for XMAS export
XMASdf <- RRI_TxHu_cleaved_full %>% filter(Confidence.Q >= 3) %>% #just get FDR<10% 
                      mutate(Crosslinks.Peptide.Group.ID = sample(100000:999999, n(), replace = FALSE),
                              "Modifications A" = "DSBSO",
                              "Modifications B" = "DSBSO",
                              "Protein Descriptions A" = "Protein.Descriptions.A", 
                              "Protein Descriptions B" = "Protein.Descriptions.B",
                              "Is Decoy" = FALSE
                              )%>%
                      select(
                      Crosslinks.Peptide.Group.ID, 
                      Crosslinks = 1,
                      Checked = 1,
                      `Crosslink Sequence` = 1, 
                      `Max. XlinkX Score` = 0, 
                      Crosslinker = 1, 
                      `Crosslink Type` = Crosslink.Type, 
                      "# CSMs" = Number.of.CSMs, 
                      "# Proteins" = 2,
                      "Sequence A" = Sequence.A, 
                      "Modifications A", 
                      "Accession A" = Accession.A, 
                      "Position A" = Position.A, 
                      "Sequence B" = Sequence.B, 
                      "Modifications B", 
                      "Accession B" = Accession.B, 
                      "Position B" = Position.B, 
                      "Protein Descriptions A", 
                      "Protein Descriptions B", 
                      "Is Decoy",
                      "Q-value" = 0,
                      "# CSMs_1" = Number.of.CSMs)
#write csv file without using "," because it is in the gene description column
write.table(XMASdf, "TachyAll_XL_publication_20240916_XMAS.csv", sep = "\t", row.names = FALSE, quote = FALSE)

#select inter and intra
inter_df <- XMASdf %>% filter(str_detect(`Crosslink Type`, "Inter"))
intra_df <- XMASdf %>% filter(str_detect(`Crosslink Type`, "Intra"))
#write csv file without using "," because it is in the gene description column
write.table(inter_df, "TachyAll_XL_publication_20240916_XMAS_inter.csv", sep = "\t", row.names = FALSE, quote = FALSE)
write.table(intra_df, "TachyAll_XL_publication_20240916_XMAS_intra.csv", sep = "\t", row.names = FALSE, quote = FALSE)
```

#Intra and Inter links distance mapping graphs
#Analyzing the distance map of intra links
```{r}
#import the tsv file that contains distance measurements.
intra_tsvdf <- read.table("TachyAll_XL_publication_20240916_XMAS_intra_LRibosome.tsv",
                      header = TRUE, 
                      fill = TRUE, 
                      sep = "\t", 
                      comment.char = "") %>% 
                  na.omit() %>%
                  mutate(Crosslink.Type = "intra")
#Small subunit
intra_tsvdf2 <- read.table("TachyAll_XL_publication_20240916_XMAS_intra_SRibosome.tsv",
                      header = TRUE, 
                      fill = TRUE, 
                      sep = "\t", 
                      comment.char = "") %>% 
                  na.omit() %>%
                  mutate(Crosslink.Type = "intra")
#combine both intralinks
intra_tsvdf <- rbind(intra_tsvdf, intra_tsvdf2)

#import the tsv file that contains distance measurements.
inter_tsvdf <- read.table("TachyAll_XL_publication_20240916_XMAS_inter_LRibosome.tsv",
                      header = TRUE, 
                      fill = TRUE, 
                      sep = "\t", 
                      comment.char = "") %>% 
                  na.omit() %>%
                  mutate(Crosslink.Type = "inter")
                  
#import the tsv file that contains distance measurements.
inter_tsvdf2 <- read.table("TachyAll_XL_publication_20240916_XMAS_inter_SRibosome.tsv",
                      header = TRUE, 
                      fill = TRUE, 
                      sep = "\t", 
                      comment.char = "") %>% 
                  na.omit() %>%
                  mutate(Crosslink.Type = "inter")

#combine both intralinks
inter_tsvdf <- rbind(inter_tsvdf, inter_tsvdf2)

# Combine "inter_tsvdf" and "intra_tsvdf" by rows
distance_df <- rbind(inter_tsvdf, intra_tsvdf)%>%
  select(Distance..A., Crosslink.Type)

#import the txt file that contains distance measurement of all possible combination of KSTY measured by Pymol
# Load all possible INTER links
inter_possible <- read.table(
  "XMAS_interprotein_ca_distancesKSTY.txt",
  header = TRUE,
  sep = "\t",
  fill = TRUE,
  fileEncoding = "Windows-1252",  # Pymol script spits out this encode!!!! Otherwise fail to load up.
  stringsAsFactors = FALSE
  )%>%
  mutate(Crosslink.Type = "possible")%>%
  rename(Distance..A. = Distance..Å.)%>%
  select(Distance..A., Crosslink.Type)

# Load all possible INTRA links
intra_possible <- read.table(
  "XMAS_intra_ca_distancesKSTY.txt",
  header = TRUE,
  sep = "\t",
  fill = TRUE,
  fileEncoding = "Windows-1252",  # Pymol script spits out this encode!!!! Otherwise fail to load up.
  stringsAsFactors = FALSE
  )%>%
  mutate(Crosslink.Type = "possible")%>%
  rename(Distance..A. = Distance..Å.)%>%
  select(Distance..A., Crosslink.Type)

# Load possible inter and intra to the 
distance_df <- bind_rows(distance_df, inter_possible, intra_possible)

# Calculate statistics for each Crosslink.Type group
grouped_summary <- distance_df %>%
  group_by(Crosslink.Type) %>%
  summarise(
    Observations = n(),
    Mean = mean(`Distance..A.`, na.rm = TRUE),
    Median = median(`Distance..A.`, na.rm = TRUE),
    SD = sd(`Distance..A.`, na.rm = TRUE)
  ) %>%
  ungroup() # Ungroup to prevent grouping affecting further operations

# Calculate overall statistics
overall_summary <- distance_df %>%
  summarise(
    Crosslink.Type = "Overall",
    Observations = n(),
    Mean = mean(`Distance..A.`, na.rm = TRUE),
    Median = median(`Distance..A.`, na.rm = TRUE),
    SD = sd(`Distance..A.`, na.rm = TRUE)
  )

# Combine the overall and grouped summaries
df_summary <- bind_rows(overall_summary, grouped_summary)
df_summary
```


#ggplot to inter and intra histogram
```{r}
library(ggplot2)
p <- ggplot(distance_df, aes(x = `Distance..A.`, fill = Crosslink.Type)) +
  geom_density(binwidth = 100, alpha = 0.6, position = "identity") + # Adjust binwidth as necessary
  scale_fill_manual(
    values = c(
      "inter" = "red", 
      "intra" = "blue", 
      "possible" = "green"
    )
  ) +
  labs(title = "Frequency Distribution of Distance by Crosslink Type",
       x = "Distance (A)", 
       y = "Frequency")
p
```
# Export the graph as an SVG file
```{r}
ggsave("XMAS_frequency_distribution_distance_by_crosslink_type.svg", plot = p, device = "svg", width = 6, height = 4)
```
#calculate the overlap between cleaved and noncleaved RRI
```{r}
# Define FDR levels and their corresponding filter conditions
fdr_levels <- list(
  FDR05 = "Confidence.Q >= 4",
  FDR01 = "Confidence.Q == 5"
)

# Function to compute counts for a given filter condition
compute_counts <- function(df_noncleaved, df_cleaved, filter_condition) {
  
  # Filter the data based on the provided condition
  RRI_noncleaved_filtered <- df_noncleaved %>% filter(eval(parse(text = filter_condition)))
  RRI_cleaved_filtered <- df_cleaved %>% filter(eval(parse(text = filter_condition)))
  
  # Total number of observations
  total_noncleaved <- RRI_noncleaved_filtered %>% summarise(count = n()) %>% pull(count)
  total_cleaved <- RRI_cleaved_filtered %>% summarise(count = n()) %>% pull(count)
  
  # Total unique RRIs
  unique_noncleaved <- RRI_noncleaved_filtered %>% distinct(RRI) %>% summarise(count = n()) %>% pull(count)
  unique_cleaved <- RRI_cleaved_filtered %>% distinct(RRI) %>% summarise(count = n()) %>% pull(count)
  
  # Shared RRIs
  shared_rris <- RRI_noncleaved_filtered %>% inner_join(RRI_cleaved_filtered, by = "RRI") %>% distinct(RRI)
  count_shared <- shared_rris %>% summarise(count = n()) %>% pull(count)
  
  # Unique RRIs only in each data frame
  unique_only_noncleaved <- unique_noncleaved - count_shared
  unique_only_cleaved <- unique_cleaved - count_shared
  
  # **New Step:** Count Crosslink Types within Shared RRIs
  # Assuming "Crosslink.Type" is present in both data frames and is consistent between them
  # We'll retrieve it from the noncleaved data frame; alternatively, you can choose one
  shared_with_type <- shared_rris %>%
    inner_join(RRI_noncleaved_filtered, by = "RRI") %>%
    select(RRI, Crosslink.Type)
  
  # Count the number of Intra and Inter types
  count_intra <- shared_with_type %>%
    filter(Crosslink.Type == "Intra") %>%
    summarise(count = n()) %>%
    pull(count)
  
  count_inter <- shared_with_type %>%
    filter(Crosslink.Type == "Inter") %>%
    summarise(count = n()) %>% 
    pull(count)
  
  # Handle cases where there might be no Intra or Inter types
  count_intra <- ifelse(length(count_intra) == 0, 0, count_intra)
  count_inter <- ifelse(length(count_inter) == 0, 0, count_inter)
  
  # Return a named vector of counts, including the new metrics
  c(
    Total_Observations_RRI_noncleaved = total_noncleaved,
    Total_Observations_RRI_cleaved = total_cleaved,
    Unique_RRIs_RRI_noncleaved = unique_noncleaved,
    Unique_RRIs_RRI_cleaved = unique_cleaved,
    Shared_RRIs = count_shared,
    Shared_RRIs_Intra = count_intra,
    Shared_RRIs_Inter = count_inter,
    Unique_RRIs_only_RRI_noncleaved = unique_only_noncleaved,
    Unique_RRIs_only_RRI_cleaved = unique_only_cleaved
  )
}

# Initialize an empty list to store results for each FDR level
results_list <- list()

# Iterate over each FDR level and compute counts
for (fdr in names(fdr_levels)) {
  counts <- compute_counts(RRI_noncleaved_full, RRI_cleaved_full, fdr_levels[[fdr]])
  results_list[[fdr]] <- counts
}

# Convert the list to a data frame
summary_df <- bind_rows(results_list, .id = "FDR") %>%
  pivot_longer(
    cols = -FDR,
    names_to = "Metric",
    values_to = "Count"
  ) %>%
  pivot_wider(
    names_from = FDR,
    values_from = Count
  )

# Optional: Arrange the metrics in a specific order
metric_order <- c(
  "Total_Observations_RRI_noncleaved",
  "Total_Observations_RRI_cleaved",
  "Unique_RRIs_RRI_noncleaved",
  "Unique_RRIs_RRI_cleaved",
  "Shared_RRIs",
  "Shared_RRIs_Intra",
  "Shared_RRIs_Inter",
  "Unique_RRIs_only_RRI_noncleaved",
  "Unique_RRIs_only_RRI_cleaved"
)

summary_df <- summary_df %>%
  mutate(Metric = factor(Metric, levels = metric_order)) %>%
  arrange(Metric)

# Replace underscores with spaces for better readability
summary_df$Metric <- gsub("_", " ", summary_df$Metric)

# (Optional) Rename Metrics for Clarity
summary_df$Metric <- case_when(
  summary_df$Metric == "Total Observations RRI noncleaved" ~ "Total Observations (noncleaved)",
  summary_df$Metric == "Total Observations RRI cleaved" ~ "Total Observations (Cleaved)",
  summary_df$Metric == "Unique RRIs RRI noncleaved" ~ "Unique RRIs (noncleaved)",
  summary_df$Metric == "Unique RRIs RRI cleaved" ~ "Unique RRIs (Cleaved)",
  summary_df$Metric == "Shared RRIs" ~ "Shared RRIs",
  summary_df$Metric == "Shared RRIs Intra" ~ "Shared RRIs (Intra)",
  summary_df$Metric == "Shared RRIs Inter" ~ "Shared RRIs (Inter)",
  summary_df$Metric == "Unique RRIs only RRI noncleaved" ~ "Unique RRIs Only (noncleaved)",
  summary_df$Metric == "Unique RRIs only RRI cleaved" ~ "Unique RRIs Only (Cleaved)",
  TRUE ~ summary_df$Metric
)

# Display the summary table
summary_df
```

#condense into PPI by grouping by PPI, count the unique R-R, then count the total spectrum
#calculate number of unique intralinks before condensing for nural network
```{r}
#before condensing, 
#calculate number of unique intralinks for each protein at FDR <10% to use for feature
intracalc <- RRI_noncleaved_FDR10 %>%
    filter(Crosslink.Type == "Intra") %>%
    mutate(Accession.A = str_remove_all(Accession.A, "\\s*\\([^)]*\\)"))%>% #remove residue info in ()
  group_by(Accession.A) %>%
  mutate(number_unique_intra = n()) %>%
  ungroup()%>%  # Optionally, remove the grouping information
  group_by(Accession.A) %>%
    # Select the row with the highest 'Best.CSM.Score' for each 'PPI'
    filter(Score == max(Score)) %>%
    ungroup() %>%
    distinct(Accession.A, .keep_all = TRUE)%>% #remove redundancy
    select(Accession.A, number_unique_intra) #select ID and interlink number

#Create a function process_PPI that encapsulates the data manipulation steps.
#This function takes two arguments:
#data: The input data frame (e.g., RRI_noncleaved_FDR10)
#confidence_q: The desired Confidence.Q value (e.g., 3)
process_PPI <- function(data, confidence_q) {
  data %>%
    group_by(PPI) %>%
    mutate(
      number_unique_RR = n(), # Count the unique Residue to Residue interactions
      Total.CSMs = sum(Number.of.CSMs, na.rm = TRUE), # Sum all CSMs within the group
      #create crosslinks for XlinkCyNET
      crosslinks_ab = paste(Accession.A, In.protein.A, Accession.B, In.protein.B, sep = "-", collapse = "#"),
      score_ab = paste(Score, collapse = "#")
    ) %>%
    filter(Score == max(Score, na.rm = TRUE)) %>% # Keep only the highest Score per PPI
    ungroup() %>%
    mutate(Best.CSM.Score = Score) %>% # Rename Score to Best.CSM.Score
    select(-c(
      Unique_Peptide_Number,
      Number.of.CSMs,
      Protein,
      In.protein.A,
      In.protein.B,
      Sequence.A,
      Sequence.B,
      Position.A,
      Position.B,
      Score
    )) %>%
    # Format for Cytoscape
    mutate(
      Accession.A = str_remove_all(Accession.A, "\\s*\\([^)]*\\)"), # Remove parenthetical content from Accession.A
      Accession.B = str_remove_all(Accession.B, "\\s*\\([^)]*\\)"), # Remove parenthetical content from Accession.B
      Confidence.Q = confidence_q # Assign the specified Confidence.Q value
    )
}

#Apply the Function to Each Dataset
#Use the process_PPI function to process each of your datasets (FDR10, FDR05, and FRD01) with their respective Confidence.Q values.

# Process FDR10 with Confidence.Q = 3
PPI_condensed_FDR10 <- process_PPI(RRI_noncleaved_FDR10, 3)
# Process FDR05 with Confidence.Q = 4
PPI_condensed_FDR05 <- process_PPI(RRI_noncleaved_FDR05, 4)
# Process FRD01 with Confidence.Q = 5
PPI_condensed_FDR01 <- process_PPI(RRI_noncleaved_FDR01, 5)

#After making PPI_condensed_FDR10, 05, 01, merge them
#merge PPI with FDR 1, 5, 10 %.
PPI_noncleaved_full <- PPI_condensed_FDR10 %>%
  mutate(Confidence.Q = if_else(PPI %in% PPI_condensed_FDR05$PPI, 4, Confidence.Q)) %>%
  mutate(Confidence.Q = if_else(PPI %in% PPI_condensed_FDR01$PPI, 5, Confidence.Q)) %>%
  filter(Crosslink.Type == "Inter") #get rid of intralinks
  
#append unique intralinks (keep the smaller of two to represent the uncertinity of the peptide)
PPI_noncleaved_full <- PPI_noncleaved_full %>%
  left_join(intracalc, by = "Accession.A") %>%
  left_join(intracalc, by = c("Accession.B" = "Accession.A"))%>%
  # Replace NA with 0 and compute the minimum of 'number_unique_intra.x' and 'number_unique_intra.y'
  mutate(number_unique_intra.min = pmin(replace_na(number_unique_intra.x, 0), replace_na(number_unique_intra.y, 0), na.rm = TRUE)) %>%
  mutate(number_unique_intra.hm = 2 * (number_unique_intra.x * number_unique_intra.y)/(number_unique_intra.x + number_unique_intra.y)) %>%
  # Remove the old columns 'number_unique_intra.x' and 'number_unique_intra.y'
  select(-number_unique_intra.x, -number_unique_intra.y)
```

#compare non-cleaved and cleaved PPIs
```{r}
#this is the cleaved PPI table
PPI_cleaved_full <- read.table("TachyAll_XL_publication_20240916_Cytoscape.Normal.csv", header = TRUE)

#set the FDR 
PPI_cleaved <- PPI_cleaved_full %>% filter(Confidence.Q >= 5)
PPI_noncleaved <- PPI_noncleaved_full %>% filter(Confidence.Q >=5) 

# Total number of observations
total_noncleaved <- PPI_noncleaved %>% summarise(count = n()) %>% pull(count)
total_cleaved <- PPI_cleaved %>% summarise(count = n()) %>% pull(count)

# Total unique PPIs
unique_noncleaved <- PPI_noncleaved %>% distinct(PPI) %>% summarise(count = n()) %>% pull(count)
unique_cleaved <- PPI_cleaved %>% distinct(PPI) %>% summarise(count = n()) %>% pull(count)

# Shared PPIs
shared_ppis <- PPI_noncleaved %>% inner_join(PPI_cleaved, by = "PPI") %>% distinct(PPI)
count_shared <- shared_ppis %>% summarise(count = n()) %>% pull(count)

# Unique PPIs only in each data frame
unique_only_noncleaved <- unique_noncleaved - count_shared
unique_only_cleaved <- unique_cleaved - count_shared

# Create summary data frame
summary_df <- tibble(
  Metric = c(
    "Total Observations in PPI_noncleaved",
    "Total Observations in PPI_cleaved",
    "Unique PPIs in PPI_noncleaved",
    "Unique PPIs in PPI_cleaved",
    "Shared PPIs between PPI_noncleaved and PPI_cleaved",
    "Unique PPIs only in PPI_noncleaved",
    "Unique PPIs only in PPI_cleaved"
  ),
  Count = c(
    total_noncleaved,
    total_cleaved,
    unique_noncleaved,
    unique_cleaved,
    count_shared,
    unique_only_noncleaved,
    unique_only_cleaved
  )
)

# View the summary data frame
print(summary_df)
```

```{r}
#export for Cytoscape
write.table(PPI_noncleaved_full, "TachyAll_XL_publication_20240916_Cytoscape.noncleaved.csv", sep = "\t", row.names = FALSE, quote = FALSE)
```
###############################################################
#
#this is true merged data for Machine Learning / Cytoscape!
#concatanate full RRI and Merge the cleaved and noncleaved PPI together for XlinkCyNET and Machine larning
```{r}

#calculate number of unique intralinks for each protein to use for feature
intracalc_full <- RRI_full %>%
    filter(Crosslink.Type == "Intra") %>%
    mutate(Accession.A = str_remove_all(Accession.A, "\\s*\\([^)]*\\)"))%>% #remove residue info in ()
  group_by(Accession.A) %>%
  mutate(number_unique_intra = n()) %>%
  ungroup()%>%  # Optionally, remove the grouping information
  group_by(Accession.A) %>%
    # Select the row with the highest 'confidence.Q' for each 'PPI'
    filter(Confidence.Q == max(Confidence.Q)) %>%
    ungroup() %>%
    distinct(Accession.A, .keep_all = TRUE)%>% #remove redundancy
    select(Accession.A, number_unique_intra) #select ID and interlink number


PPI_RRI_full <- RRI_full %>%
  group_by(PPI) %>%
    mutate(
      number_unique_RR = n(), # Count the unique Residue to Residue interactions
      Total.CSMs = sum(Number.of.CSMs, na.rm = TRUE), # Sum all CSMs within the group
      #create crosslinks for XlinkCyNET
      crosslinks_ab = paste(Accession.A, In.protein.A, Accession.B, In.protein.B, sep = "-", collapse = "#"),
      score_ab = paste(Confidence.Q, collapse = "#"),
      # Assign MS2 based on the values in MS
    MS = case_when(
      all(MS == "cleaved") ~ "cleaved",
      all(MS == "noncleaved") ~ "noncleaved",
      TRUE ~ "both"),
    Confidence.Q = max(Confidence.Q, na.rm = TRUE),
    Best.CSM.Score = max(Best.CSM.Score, na.rm = TRUE)
    ) %>% # Keep only the highest Score per PPI
    slice_max(order_by = Confidence.Q, n = 1, with_ties = FALSE) %>% # Retain only the top row per PPI
    ungroup() %>%
  
    select(-c(
      Number.of.CSMs,
      RRI,
      In.protein.A,
      In.protein.B,
      Sequence.A,
      Sequence.B,
      Position.A,
      Position.B
    )) %>%
#append unique intralinks (keep the smaller of two to represent the uncertinity of the peptide)
  left_join(intracalc, by = "Accession.A") %>%
  left_join(intracalc, by = c("Accession.B" = "Accession.A"))%>%
  # Replace NA with 0 and compute the minimum of 'number_unique_intra.x' and 'number_unique_intra.y'
  mutate(number_unique_intra.min = pmin(replace_na(number_unique_intra.x, 0), replace_na(number_unique_intra.y, 0), na.rm = TRUE))%>%
  # Remove the old columns 'number_unique_intra.x' and 'number_unique_intra.y'
  select(-number_unique_intra.x, -number_unique_intra.y)
```

#Format for Cytoscape with RRI for XlinkCyNET
```{r}
#you need the protein length for XlinkCyNET. Get them from the Gene Name Lookup Table
gene.name <- read.table(file = "GeneNameLookupTableTxHu.tsv", sep = "\t", header = TRUE, quote = "") %>%
    distinct(gene, .keep_all = TRUE)

#add protein lengths
XlinkCyNETdf <- PPI_RRI_full %>%
  # First left_join for Accession.A
  left_join(
    gene.name %>% select(gene, protein_length),      # Select only necessary columns
    by = c("Accession.A" = "gene")                 # Join keys
  ) %>%
  rename(length_protein_a = protein_length) %>%      # Rename the joined column

  # Second left_join for Accession.B
  left_join(
    gene.name %>% select(gene, protein_length),      # Select only necessary columns
    by = c("Accession.B" = "gene")                 # Join keys
  ) %>%
  rename(length_protein_b = protein_length)%>%          # Rename the joined column

  #XlinkCyNET format
  mutate(
  gene_a = Accession.A,
  gene_b = Accession.B,
  ppi_score = as.numeric(Confidence.Q) * as.numeric(number_unique_RR),
  crosslinks_ba = crosslinks_ab,
  score_ba = score_ab,
  protein_a = gene_a,
  protein_b = gene_b
)

#write the Cytoscape PPI data for XlinkCyNET
write.table(XlinkCyNETdf %>% filter(Crosslink.Type == "Inter", Confidence.Q >= 3),
  file = "TachyAll_XL_publication_20240916_XlinkCyNET.tsv",    # summary of dimers
  sep = "\t", quote = FALSE, row.names = FALSE, col.names = TRUE)

#write the Cytoscape PPI data
write.table(PPI_RRI_full %>% 
              filter(Crosslink.Type == "Inter", Confidence.Q >= 3),
  file = "TachyAll_XL_publication_20240916_Cytoscape.tsv",    # summary of dimers
  sep = "\t", quote = FALSE, row.names = FALSE, col.names = TRUE)

```

#from here use the NeuralNetwork PPI script to get the prediction
#load the features and STRING, append it, then engineer it for prediction
```{r}
#Read the features into a text file if skipping the feature set creation
featureDF <- read.table(file = "ToxoDB67_AllFeature_woSTRING.txt", sep = "\t", header = TRUE)
STRINGdf <- read.table(file = "ToxoDB67_AllSTRING.txt", sep = "\t", header = TRUE)
#append featureDF to the main XLMS data frame
df <- PPI_RRI_full %>%
  left_join(featureDF, by = c("Accession.A" = "Gene_ID"))%>%
  left_join(featureDF, by = c("Accession.B" = "Gene_ID"))%>%
  left_join(STRINGdf, by = "PPI") %>%
  rename(STRING.score = combined_score) #rename the column name score to reflect STRING
#Checking if there is any match in GO term, hyperLOPIT, etc.
#Define the Matching Function: We'll start with the matching function. It checks for any common elements between two semicolon-separated strings.

find_matches_boolean <- function(x, y) {
  # Convert inputs to character, in case they are not
  x <- as.character(x)
  y <- as.character(y)
  
  # Pre-calculate NA or "0" check
  na_or_zero_check <- is.na(x) || is.na(y) || (x == "0" && y == "0")
  
  # Split the strings into vectors based on the ";" delimiter, if not NA or "0"
  if (!na_or_zero_check) {
    items_x <- unlist(strsplit(x, split = ";", fixed = TRUE))
    items_y <- unlist(strsplit(y, split = ";", fixed = TRUE))
    
    # Return TRUE if there are any matches, taking into account the NA or "0" check
    return(length(intersect(items_x, items_y)) > 0)
  } else {
    # Return NA if either input is NA or both are "0"
    return(NA)
  }
}

#Identify and Process Column Pairs: We'll identify the pairs of columns that need to be compared and then iterate over these pairs, applying the matching function to each pair and storing the result in a new column.
# Extract column names
col_names <- names(df)

# Identify unique prefixes by removing the trailing .x and .y
prefixes <- unique(sub("\\.(x|y)$", "", col_names))

# Filter out prefixes to ensure both .x and .y versions exist
valid_prefixes <- prefixes[sapply(prefixes, function(prefix) {
  all(c(paste0(prefix, ".x"), paste0(prefix, ".y")) %in% col_names)
})]

# Iterate over each valid prefix and compare the .x and .y columns
for (prefix in valid_prefixes) {
  x_col <- paste0(prefix, ".x")
  y_col <- paste0(prefix, ".y")
  match_col <- paste0(prefix, "_matches") # New column for match results
  
  # Apply the matching function to each row for the current pair of columns
  df[[match_col]] <- mapply(find_matches_boolean, df[[x_col]], df[[y_col]], SIMPLIFY = TRUE)
}


# Define the function Q(x, y) for calculating absolute difference, 2x if they are + and -
Q <- function(x, y) {
  D <- abs(x - y)
  delta <- ifelse(sign(x) != sign(y), 1, 0)
  Q_value <- D * (1 + delta)
  return(Q_value)
}

#calculate absolute difference between CRISPR score, 2X if they are + and -
df <- df %>%
  mutate(CRISPR.x = as.numeric(CRISPR.x),
         CRISPR.y = as.numeric(CRISPR.y),
         CRISPR.dif = Q(CRISPR.x, CRISPR.y))
#calculate absolute difference between Non synonimous synonimous SNP ratio, 2X if they are + and -
df <- df %>%
  mutate(NonSyn_Syn_SNP_Ratio_All_Strains_c.x = as.numeric(NonSyn_Syn_SNP_Ratio_All_Strains_c.x),
         NonSyn_Syn_SNP_Ratio_All_Strains_c.y = as.numeric(NonSyn_Syn_SNP_Ratio_All_Strains_c.y),
        NonSyn_Syn_SNP_Ratio_All_Strains.dif = Q(NonSyn_Syn_SNP_Ratio_All_Strains_c.x, NonSyn_Syn_SNP_Ratio_All_Strains_c.y))

#calculate minimum for RNAseq # Calculate the minimum of RnaSeq.x and RnaSeq.y
df <- df %>%
  rowwise() %>%
  mutate(RnaSeq.min = min(c(RnaSeq.x, RnaSeq.y), na.rm = TRUE)) %>%
  ungroup()

#get rid of unnecessary columns
df <- df %>% select(
  PPI,
  Accession.A,
  Accession.B,
  Confidence.Q,
  MS,
  Crosslink.Type,
  number_unique_RR,
  Total.CSMs,
  number_unique_intra.min,
  #number_unique_intra.hm,
  Computed_GO_Component_IDs_c_matches,
  Computed_GO_Function_IDs_c_matches,
  Computed_GO_Process_IDs_c_matches,
  EC_numbers_from_OrthoMCL_c_matches,
  PFam_ID_c_matches,
  ToxoNET_matches,
  Location_matches,
  CRISPR.dif,
  NonSyn_Syn_SNP_Ratio_All_Strains.dif,
  RnaSeq.min,
  STRING.score,
  Best.CSM.Score
)

#Write the data frame into a text file
write.table(df %>% filter(Crosslink.Type == "Inter"), file = "TachyAll_XL_publication_20240916_PPI_combined_wFeatures.txt", sep = "\t", row.names = FALSE, col.names = TRUE)
```

###After lightGBM Predicted###################################
#Prediction with combined did not work well. Just used the cleaved model to predict combined

```{r}
raw_pred <- read.table("TachyAll_XL_publication_20240916_PPI_combined_wFeatures_with_lightGBM_probabilities.tsv", header = TRUE, sep = "\t") 

PPI_pred <- raw_pred %>% select(
  Accession.A,
  Accession.B,
  Confidence.Q,
  number_unique_RR,
  Total.CSMs,
  PPI,
  MS,
  lightGBM_prob
) %>%
  filter(Confidence.Q != 1)

#Salvage the low score hits with high probability
PPI_pred_filtered <- PPI_pred %>% filter(Confidence.Q == 5 & lightGBM_prob > 0.2 | #high FDR < 1 (Q=5)
                                   (Confidence.Q == 4 & lightGBM_prob > 0.5) | #medium FDR < 5 (Q=4)
                                   (Confidence.Q == 3 & lightGBM_prob > 0.6)| #low FDR <10 (Q=3)
                                    (Confidence.Q == 2 & lightGBM_prob > 0.9)| #purgatory FDR <30 (Q=2)
                                   (Confidence.Q == 1 & lightGBM_prob == 1) #hell FDR >30
                              )
# Count observations grouped by Confidence.Q for both data frames
filtered_counts <- PPI_pred_filtered %>% count(Confidence.Q, name = "Count") %>% mutate(Source = "filtered")
filtered_non_counts <- PPI_pred_filtered %>% 
  filter(MS == "noncleaved") %>%
  count(Confidence.Q, name = "Count") %>% mutate(Source = "filtered_non_pred")
raw_pred_counts <- raw_pred %>% count(Confidence.Q, name = "Count") %>% mutate(Source = "raw_pred")
raw_non_counts <- raw_pred %>% 
  filter(MS == "noncleaved") %>%
  count(Confidence.Q, name = "Count") %>% mutate(Source = "raw_non_pred")

# Combine into one table
combined_counts <- bind_rows(filtered_counts, raw_pred_counts, filtered_non_counts, raw_non_counts)

combined_counts %>%
  pivot_wider(names_from = Source, values_from = Count, values_fill = 0)%>%
  arrange(Confidence.Q)  # Sorting based on Confidence.Q
# Save file for Interactome Cytoscape
write.table(PPI_pred_filtered, file = "TachyAll_XL_publication_20240916_PPI_PRED_Cytoscape.tsv", sep = "\t", row.names = FALSE, col.names = TRUE, quote = FALSE)
```
#prepare the file for RRI interaction with the individual residues so that we can use XlinkCyNET
Prerequisite: PPI_RRI_full
```{r}
#append protein length and gene description from the gene neme lookup
gene.name <- read.table(file = "GeneNameLookupTableTxHu.tsv", sep = "\t", header = TRUE, quote = "") %>%
    distinct(gene, .keep_all = TRUE)
#join the crosslink RRI information from PPI_RRI_full
PPI_RRI_pred_XlinkCyNET <- left_join(PPI_pred_filtered, PPI_RRI_full %>% select(PPI,
                                                                     Best.CSM.Score,
                                                                     crosslinks_ab,
                                                                     score_ab), 
                          by = "PPI") %>%
#append protein length and gene name and rename to XlinkCyNET format
  left_join(gene.name, by = c("Accession.A" = "gene")) %>%
  left_join(gene.name, by = c("Accession.B" = 'gene')) %>%
  rename(gene_product.a = gene_product.x,
         gene_product.b = gene_product.y,
         length_protein_a = protein_length.x,
         length_protein_b = protein_length.y,
         gene_a = Accession.A,
         gene_b = Accession.B,
         ) %>%
  mutate(
  ppi_score = as.numeric(Confidence.Q) * as.numeric(Total.CSMs),
  crosslinks_ba = crosslinks_ab,
  score_ba = score_ab,
  protein_a = gene_a,
  protein_b = gene_b
        ) %>%
  select(-c(species.x, species.y))

# Save file for XlinkCyNET Cytoscape
write.table(PPI_RRI_pred_XlinkCyNET, file = "TachyAll_XL_publication_20240916_PPI_PRED_XlinkCyNET.tsv", sep = "\t", row.names = FALSE, col.names = TRUE, quote = FALSE)

```
#### Dimer detection
Prerequisit: need RRI_full that has all the residue-to-residue interaction
```{r}
dimer_df <- RRI_full %>%
  filter(Confidence.Q >= 3)%>% #FDR <5%
  filter(Accession.A == Accession.B) %>%
    #there are multiple positions due to the multiple protein mapping to a peptide sequence
  mutate(
    # Extract the first number before ";" and convert to numeric
    In.protein.A = as.numeric(str_split(In.protein.A, ";", simplify = TRUE)[,1]),
    In.protein.B = as.numeric(str_split(In.protein.B, ";", simplify = TRUE)[,1]),

    # Calculate Peptide.A.Start and Peptide.B.Start
    Peptide.A.Start = In.protein.A - Position.A + 1,
    Peptide.B.Start = In.protein.B - Position.B + 1,
    
    # Extract substring after ']' in Sequence.A and Sequence.B
    # and calculate Peptide.A.End and Peptide.B.End
    Peptide.A.End = In.protein.A + str_length(str_remove(Sequence.A, "^.*\\]")),
    Peptide.B.End = In.protein.B + str_length(str_remove(Sequence.B, "^.*\\]")),
    
    # Determine if there is any overlap between the peptide ranges
    Dimer = (Peptide.A.Start <= Peptide.B.End) & (Peptide.B.Start <= Peptide.A.End)
  ) %>%
    filter(Dimer == TRUE)

#concatenate the dimer and create number of RRI per dimerized gene, create XlinkCyNET format
dimer_detect <- dimer_df %>%
  group_by(Accession.A) %>%
  summarise(dimer_RRI = n(),
            crosslinks_ab = paste(Accession.A, In.protein.A, Accession.B, In.protein.B, sep = "-", collapse = "#"),
            score_ab = paste(Confidence.Q, collapse = "#")
            ) %>%
  ungroup() 

gene.name <- read.table(file = "GeneNameLookupTableTxHu.csv", sep = "\t", header = TRUE, quote = "") %>%
    distinct(gene, .keep_all = TRUE)

#add gene description to the summary
dimer_detect <- dimer_detect %>%
  left_join(gene.name, by = c("Accession.A" = "gene")) %>%
  #XlinkCyNET format
  mutate(
  gene_a = Accession.A,
  gene_b = Accession.A,
  ppi_score = as.numeric(score_ab) * as.numeric(dimer_RRI),
  crosslinks_ba = crosslinks_ab,
  score_ba = score_ab,
  protein_a = gene_a,
  protein_b = gene_b
)

#add gene description to the dimer data frame
dimer_df <- dimer_df %>%
  left_join(gene.name, by = c("Accession.A" = "gene"))

#write the summary of dimer
write.table(dimer_detect,
  file = "dimer_detect.tsv",    # summary of dimers
  sep = "\t", quote = FALSE, row.names = FALSE, col.names = TRUE)

#write actual dimer RRI
write.table(dimer_df,
  file = "dimer_df.tsv",    # this is detailed RRI for dimers
  sep = "\t", quote = FALSE, row.names = FALSE, col.names = TRUE)
```


##### Machine Learning with Python ####
import lightgbm as lgb
import pandas as pd
import numpy as np
import optuna # Import Optuna for hyperparameter optimization
import logging
from sklearn.model_selection import StratifiedKFold, train_test_split
from sklearn.metrics import (
    accuracy_score,
    roc_auc_score,
    confusion_matrix,
    classification_report,
)
import matplotlib.pyplot as plt
import seaborn as sns
import shap
from datetime import datetime
import os
import re  # Ensure regex module is imported for sanitize_filename

# -----------------------------------------------
# Minimal Logging Setup
# -----------------------------------------------
def setup_logging(log_file='lightgbm_workflow.log'):
    logger = logging.getLogger()
    logger.setLevel(logging.INFO)

    # Clear existing handlers to prevent duplication
    if logger.hasHandlers():
        logger.handlers.clear()

    # Create handlers
    file_handler = logging.FileHandler(log_file)
    console_handler = logging.StreamHandler()

    # Set levels
    file_handler.setLevel(logging.INFO)
    console_handler.setLevel(logging.INFO)

    # Create formatters and add to handlers
    formatter = logging.Formatter("%(asctime)s [%(levelname)s] %(message)s")
    file_handler.setFormatter(formatter)
    console_handler.setFormatter(formatter)

    # Add handlers to the logger
    logger.addHandler(file_handler)
    logger.addHandler(console_handler)

    return logger

# Initialize logger
logger = setup_logging()

# Example log messages
logger.info("Logging has been set up successfully.")

# -----------------------------------------------
# Define the sanitize_filename Function
# -----------------------------------------------
def sanitize_filename(name):
    """
    Sanitize the feature name to create a safe filename by replacing
    non-alphanumeric characters with underscores.

    Parameters:
    - name (str): The original feature name.

    Returns:
    - str: The sanitized feature name.
    """
    # Replace any character that is not alphanumeric or underscore with an underscore
    return re.sub(r'[^A-Za-z0-9_]+', '_', name)

# -----------------------------------------------
# 4. Load the Dataset
# -----------------------------------------------
# If using Google Colab, mount Google Drive
try:
    from google.colab import drive
    drive.mount('/content/drive')  # This mounts your Google Drive
    data_path = '/content/drive/MyDrive/Programming/Colab_XLMS_MachineLearning/TachyAll_XL_publication_20240916_PPI_wFeatures.txt'
except:
    # If not using Colab, specify the local path
    data_path = 'path_to_your_local_data/TachyAll_XL_publication_20240916_PPI_wFeatures.txt'

# Extract data directory and filename
data_dir = os.path.dirname(data_path)
data_filename = os.path.basename(data_path)
data_name, data_ext = os.path.splitext(data_filename)

# Load the dataset
df = pd.read_csv(data_path, delimiter='\t')
logger.info("Dataset loaded successfully.")

# Display dataset information
logger.info(f"Dataset shape: {df.shape}")
logger.info(f"Dataset columns: {df.columns.tolist()}")

# -----------------------------------------------
# Define the Labeling Function and Apply It
# -----------------------------------------------
def label_csm_one(confidence):
    if confidence < 10:  # Negative data
        return 0
    elif confidence >= 235:  # Positive data
        return 1
    else:
        return np.nan  # Use NaN for easier filtering

df['label'] = df['Best.CSM.Score'].apply(label_csm_one)
logger.info("Labeling function applied.")

# Filter out rows with NaN labels (all mediocre hits)
data_filtered = df[df['label'].notnull()].copy()
logger.info(f"Filtered dataset shape (after labeling): {data_filtered.shape}")

# -----------------------------------------------
# 6. Feature Selection
# -----------------------------------------------
features = [
    'Location_matches',
    'STRING.score',
    'RnaSeq.min',
    'ToxoNET_matches',
    'Computed_GO_Component_IDs_c_matches',
    'Computed_GO_Function_IDs_c_matches',
    'Computed_GO_Process_IDs_c_matches',
    'EC_numbers_from_OrthoMCL_c_matches',
    'PFam_ID_c_matches',
    'CRISPR.dif',
    'NonSyn_Syn_SNP_Ratio_All_Strains.dif',
    'Total.CSMs',
    'number_unique_RR',
    'number_unique_intra.min'
    #'number_unique_intra.hm'
]

# Ensure selected features exist in the dataset
missing_features = set(features) - set(data_filtered.columns)
if missing_features:
    logger.error(f"Missing features in the dataset: {missing_features}")
    raise ValueError(f"Missing features: {missing_features}")

X = data_filtered[features]
y = data_filtered['label'].astype(int)
logger.info("Features and labels extracted.")

# -----------------------------------------------
# Data Preprocessing: Convert Boolean Features
# -----------------------------------------------
# List of features with TRUE, FALSE, NA values
boolean_features = [
    'Location_matches',
    'ToxoNET_matches',
    'Computed_GO_Component_IDs_c_matches',
    'Computed_GO_Function_IDs_c_matches',
    'Computed_GO_Process_IDs_c_matches',
    'EC_numbers_from_OrthoMCL_c_matches',
    'PFam_ID_c_matches'
]

# Replace 'NA' strings with np.nan in the entire dataset
X.replace('NA', np.nan, inplace=True)

# Diagnose: Check unique values before conversion
logger.info("Checking unique values in boolean features before conversion:")
for col in boolean_features:
    unique_values = X[col].unique()
    logger.info(f"Unique values in '{col}': {unique_values}")

# Convert boolean features to numeric
for col in boolean_features:
    # Strip whitespace and convert to uppercase
    X[col] = X[col].astype(str).str.strip().str.upper()

    # Map actual unique values to 1/0
    mapping_dict = {'TRUE': 1, 'FALSE': 0}
    X[col] = X[col].map(mapping_dict)

    # Ensure the column is of float dtype to accommodate np.nan
    X[col] = X[col].astype(float)

logger.info("Boolean features converted to numeric.")

# Display a summary to check the conversion
logger.info("Summary of boolean features after conversion:")
print(X[boolean_features].describe())

# Display counts of unique values after conversion
for col in boolean_features:
    value_counts = X[col].value_counts(dropna=False)
    logger.info(f"Value counts for '{col}' after conversion:\n{value_counts}")


# -----------------------------------------------
# Split the Data into Training and Test Sets
# -----------------------------------------------
# Split data with stratification to maintain class proportions
X_train, X_test, y_train, y_test = train_test_split(
    X, y,
    test_size=0.1,  # Set aside 10% for testing
    stratify=y,
    random_state=42
)
logger.info(f"Training set shape: {X_train.shape}")
logger.info(f"Test set shape: {X_test.shape}")

# -----------------------------------------------
# Verify Data Types After Conversion
# -----------------------------------------------
logger.info("Data types in training set after conversion:")
logger.info(X_train.dtypes)

# -----------------------------------------------
# 8. Prepare LightGBM Dataset for Cross-Validation
# -----------------------------------------------
lgb_dataset = lgb.Dataset(
    X_train,
    label=y_train,
    free_raw_data=False
)

# -----------------------------------------------
# 9. Define Model Parameters
# -----------------------------------------------
params = {
    'objective': 'binary',
    'metric': ['auc', 'binary_logloss'],  # 'auc' first for early stopping
    'boosting_type': 'gbdt',
    #'num_leaves': 31, it will be tuned by optuna
    #'learning_rate': 0.05, it will be tuned by optuna
    'feature_fraction': 0.9,
    'bagging_fraction': 0.8,
    'bagging_freq': 5,
    'verbosity': -1,
    'random_state': 42,
    'is_unbalance': True  # Set to True for class imbalance handling
}

# -----------------------------------------------
# 10. Hyperparameter Tuning with Optuna
# -----------------------------------------------
logger.info("Starting hyperparameter tuning with Optuna.")

def objective(trial):
    # Define the hyperparameter search space
    param = {
        'objective': 'binary',
        'metric': 'auc',
        'boosting_type': 'gbdt',
        'is_unbalance': True,
        'random_state': 42,
        'num_leaves': trial.suggest_int('num_leaves', 15, 127),
        'learning_rate': trial.suggest_float('learning_rate', 0.005, 0.3, log=True),
        'max_depth': trial.suggest_int('max_depth', 5, 50),
        'min_child_samples': trial.suggest_int('min_child_samples', 5, 100),
        'subsample': trial.suggest_float('subsample', 0.5, 1.0),
        'colsample_bytree': trial.suggest_float('colsample_bytree', 0.5, 1.0),
        'reg_alpha': trial.suggest_float('reg_alpha', 0.0, 10.0),
        'reg_lambda': trial.suggest_float('reg_lambda', 0.0, 10.0),
    }


# -----------------------------------------------
# 11. Set Up Cross-Validation with Callbacks
# -----------------------------------------------
    n_splits = 3
    skf = StratifiedKFold(n_splits=n_splits, shuffle=True, random_state=49)

    logger.info(f"Starting {n_splits}-Fold Stratified Cross-Validation.")

# Define callbacks for cross-validation
# you need this for Early Stopping.
    callbacks_cv = [
    lgb.log_evaluation(period=10),  # Logs every 10 rounds
    lgb.early_stopping(stopping_rounds=100, verbose=True)
    ]

# Perform cross-validation with callbacks
    cv_results = lgb.cv(
      param, #changed to param from the params !!!
      lgb_dataset,
      num_boost_round=1000,
      folds=skf.split(X_train, y_train),
      seed=49,
      stratified=True, #do we need this?
      callbacks=callbacks_cv
    )

    # Extract the best score
    best_score = max(cv_results['valid auc-mean']) #auc-mean is valid auc-mean
    return best_score

# Create a study object and optimize the objective function
study = optuna.create_study(direction='maximize', study_name='lightgbm_hyperparameter_tuning')
study.optimize(objective, n_trials=100)

# Log the best hyperparameters
logger.info(f"Best hyperparameters found by Optuna: {study.best_params}")
logger.info(f"Best AUC from Optuna: {study.best_value:.4f}")

# Update the parameters with the best hyperparameters
params.update(study.best_params)

# Since 'metric' might be overwritten, ensure it's included
params['metric'] = ['auc', 'binary_logloss']


# -----------------------------------------------
# 12. Train Final Model on Full Training Set with Early Stopping
# -----------------------------------------------
logger.info("Training final model with early stopping on validation set using best hyperparameters.")

# Split X_train into final training and validation sets
X_train_final, X_val, y_train_final, y_val = train_test_split(
    X_train,
    y_train,
    test_size=0.1,  # Allocate 10% for validation
    stratify=y_train,
    random_state=49
)
logger.info(f"Final Training set shape: {X_train_final.shape}")
logger.info(f"Validation set shape: {X_val.shape}")

# Create LightGBM Datasets
lgb_train_final = lgb.Dataset(
    X_train_final,
    label=y_train_final,
    free_raw_data=False
)

lgb_val = lgb.Dataset(
    X_val,
    label=y_val,
    reference=lgb_train_final,
    free_raw_data=False
)

# Define callbacks for final training
callbacks_train = [
    lgb.log_evaluation(period=50),  # Logs every 50 rounds
    lgb.early_stopping(stopping_rounds=100, verbose=True)
]

# Train the final model with early stopping
final_model = lgb.train(
    params,
    lgb_train_final,
    num_boost_round=1000,
    valid_sets=[lgb_val],
    valid_names=['validation'],
    callbacks=callbacks_train
)

logger.info("Final model trained successfully.")
logger.info(f"Final model best iteration: {final_model.best_iteration}")

# -----------------------------------------------
# Evaluate the Model on the Test Set
# -----------------------------------------------
logger.info("Evaluating the model on the test set.")

# Predict probabilities on the test set
y_pred_proba = final_model.predict(X_test, num_iteration=final_model.best_iteration)

# For binary classification, set a threshold of 0.5
y_pred = (y_pred_proba >= 0.5).astype(int)

# Compute performance metrics
accuracy = accuracy_score(y_test, y_pred)
roc_auc = roc_auc_score(y_test, y_pred_proba)
conf_matrix = confusion_matrix(y_test, y_pred)
classif_report = classification_report(y_test, y_pred)

# Log the performance metrics
logger.info(f"Test Set Accuracy: {accuracy:.4f}")
logger.info(f"Test Set ROC AUC: {roc_auc:.4f}")
logger.info("Confusion Matrix:")
logger.info(f"\n{conf_matrix}")
logger.info("Classification Report:")
logger.info(f"\n{classif_report}")

# Optionally, plot the ROC curve
from sklearn.metrics import roc_curve, auc

fpr, tpr, thresholds = roc_curve(y_test, y_pred_proba)
roc_auc_curve = auc(fpr, tpr)

# Plot ROC curve
plt.figure()
plt.plot(fpr, tpr, color='darkorange', lw=2, label=f'ROC curve (area = {roc_auc_curve:.4f})')
plt.plot([0, 1], [0, 1], color='navy', lw=2, linestyle='--')
plt.xlim([-0.05, 1.0])
plt.ylim([0.0, 1.05])
plt.xlabel('False Positive Rate')
plt.ylabel('True Positive Rate')
plt.title('Receiver Operating Characteristic - Test Set')
plt.legend(loc="lower right")

# Save the ROC curve as SVG
roc_curve_filename = f"{data_name}_ROC_Curve.svg"
roc_curve_path = os.path.join(data_dir, roc_curve_filename)
plt.savefig(roc_curve_path, format='svg')
plt.close()
logger.info(f"ROC curve saved to {roc_curve_path}")

# -----------------------------------------------
# 13. Feature Importance
# -----------------------------------------------
# Get feature importances
importance = final_model.feature_importance(importance_type='split')  # or 'gain'
feature_names = final_model.feature_name()

# Create a DataFrame for visualization
feature_importance_df = pd.DataFrame({
    'feature': feature_names,
    'importance': importance
})

# Sort the DataFrame
feature_importance_df = feature_importance_df.sort_values(by='importance', ascending=False)

# Plot
plt.figure(figsize=(10, 8))
sns.barplot(x='importance', y='feature', data=feature_importance_df.head(20), palette='magma')
plt.title('Top 20 Feature Importances')
plt.xlabel('Importance (split)')
plt.ylabel('Feature')
plt.tight_layout()

# Save the feature importance plot as SVG
feature_importance_filename = f"{data_name}_Feature_Importance.svg"
feature_importance_path = os.path.join(data_dir, feature_importance_filename)
plt.savefig(feature_importance_path, format='svg')
plt.close()
logger.info(f"Feature importance plot saved to {feature_importance_path}")

# -----------------------------------------------
# 14. Explainability by SHAP
# -----------------------------------------------
logger.info("Calculating SHAP values for model explainability.")

# Create a TreeExplainer for LightGBM
explainer = shap.TreeExplainer(final_model)
shap_values = explainer.shap_values(X_train_final)

# Plot SHAP summary
shap.summary_plot(shap_values, X_train_final, show=False)

# Save the SHAP summary plot as SVG
shap_summary_filename = f"{data_name}_SHAP_Summary.svg"
shap_summary_path = os.path.join(data_dir, shap_summary_filename)
plt.savefig(shap_summary_path, format='svg')
plt.close()
logger.info(f"SHAP summary plot saved to {shap_summary_path}")

# -----------------------------------------------
# Calculate Mean Absolute SHAP Values
# -----------------------------------------------
logger.info("Calculating Mean Absolute SHAP values.")

# Compute the mean absolute SHAP value for each feature
mean_abs_shap = np.abs(shap_values).mean(axis=0)
shap_importance_df = pd.DataFrame({
    'feature': X_train_final.columns,
    'mean_abs_shap': mean_abs_shap
}).sort_values(by='mean_abs_shap', ascending=False)

# Log the Mean Absolute SHAP values
logger.info("Mean Absolute SHAP values:")
for index, row in shap_importance_df.iterrows():
    logger.info(f"{row['feature']}: {row['mean_abs_shap']:.4f}")

# Plot Mean Absolute SHAP values
plt.figure(figsize=(10, 8))
sns.barplot(x='mean_abs_shap', y='feature', data=shap_importance_df.head(20), palette='viridis')
plt.title('Top 20 Features by Mean Absolute SHAP Values')
plt.xlabel('Mean |SHAP value|')
plt.ylabel('Feature')
plt.tight_layout()

# Save the Mean Absolute SHAP importance plot as SVG
shap_importance_filename = f"{data_name}_Mean_Absolute_SHAP_Importance.svg"
shap_importance_path = os.path.join(data_dir, shap_importance_filename)
plt.savefig(shap_importance_path, format='svg')
plt.close()
logger.info(f"Mean Absolute SHAP importance plot saved to {shap_importance_path}")

# -----------------------------------------------
# Generate and Save SHAP Dependence Plots
# -----------------------------------------------
logger.info("Generating SHAP dependence plots for each feature.")

# Create a subdirectory for SHAP dependence plots
shap_dependence_dir = os.path.join(data_dir, f"{data_name}_SHAP_Dependence_Plots")
os.makedirs(shap_dependence_dir, exist_ok=True)

for feature in X_train_final.columns:
    plt.figure()
    shap.dependence_plot(
        feature,
        shap_values,
        X_train_final,
        show=False,
        interaction_index=None  # Set to None to not color by any feature
    )

    # Define the filename by appending the sanitized feature name to the data file name
    sanitized_feature = sanitize_filename(feature)
    shap_dependence_filename = f"{data_name}_SHAP_Dependence_{sanitized_feature}.svg"
    shap_dependence_path = os.path.join(shap_dependence_dir, shap_dependence_filename)

    # Save the SHAP dependence plot as SVG
    plt.savefig(shap_dependence_path, format='svg')
    plt.close()
    logger.info(f"SHAP dependence plot for '{feature}' saved to {shap_dependence_path}")

# -----------------------------------------------
# 15. Save the Final Model to a File
# -----------------------------------------------
# Construct the new model filename
model_filename = f"{data_name}_model.txt"  # Using '.txt' as the file extension
model_path = os.path.join(data_dir, model_filename)

# Save the final model to the model_path
final_model.save_model(model_path)
logger.info(f"Final model saved to {model_path}")

# -----------------------------------------------
# Use the Trained Model to Predict on the Full Dataset
# -----------------------------------------------
logger.info("Loading the original dataset for full prediction.")

# Load the original dataset
df_full = pd.read_csv(data_path, delimiter='\t')
logger.info("Original dataset loaded successfully.")

# Extract features
X_full = df_full[features]
# Replace 'NA' strings with np.nan in the entire dataset
X_full.replace('NA', np.nan, inplace=True)
# Convert boolean features to numeric
for col in boolean_features:
    # Strip whitespace and convert to uppercase
    X_full[col] = X_full[col].astype(str).str.strip().str.upper()

    # Map actual unique values to 1/0
    mapping_dict = {'TRUE': 1, 'FALSE': 0}
    X_full[col] = X_full[col].map(mapping_dict)

    # Ensure the column is of float dtype to accommodate np.nan
    X_full[col] = X_full[col].astype(float)

# Display a summary to check the conversion
logger.info("Summary of boolean features after conversion:")
print(X_full[boolean_features].describe())

#use the same model to predict the probability for each observation in X_full and append it to the df_full as a column named "lightGBM_prob"

# Predict probabilities using the trained final_model
y_full_pred_proba = final_model.predict(
    X_full,
    num_iteration=final_model.best_iteration
)

# Append the predicted probabilities to df_full as a new column
df_full['lightGBM_prob'] = y_full_pred_proba
logger.info("Predicted probabilities appended to df_full as 'lightGBM_prob'.")

# Optionally, save the updated df_full to a new file
# Define the output filename
output_filename = f"{data_name}_with_lightGBM_probabilities.tsv"
output_path = os.path.join(data_dir, output_filename)
# Save as a TSV file
df_full.to_csv(output_path, sep='\t', index=False)
logger.info(f"Updated df_full with probabilities saved to {output_path}")
